# *Chlamydomonas* chloroplast genes tolerate compression of the genetic code to just 51 codons

**DOI:** 10.1101/2025.03.09.641718

**Authors:** Pawel M. Mordaka, Kitty Clouston, Jing Cui, Andre Holzer, Harry O. Jackson, Saul Purton, Alison G. Smith

## Abstract

Genome scale engineering has enabled codon compression of the universal genetic code of up to three codons in *E. coli,* providing the means for genetic code expansion. To go much beyond this number, smaller and simpler genetic systems are needed to avoid significant technical challenges. Chloroplast genomes offer multiple advantages for codon compression and reassignment. Here we report a recoding scheme for the *Chlamydomonas reinhardtii* chloroplast genome, in which two stop codons and one or more of the codons for arginine, glycine, isoleucine, leucine and serine, all of which have two cognate tRNAs, are absent, compressing the genetic code to 51 codons. Firstly, several recoding strategies were tested on the essential *rpoA* gene, encoding a subunit of the chloroplast RNA polymerase. A defined compression scheme, which relied on swapping the target codons with the permitted frequent codons, could replace the native protein coding sequence without affecting chloroplast protein expression levels or the strain fitness. The same strategy was successfully used for codon compression of *ycf1*, encoding a subunit of the chloroplast translocon, *psaA* and *psbA*, intron-containing highly expressed genes encoding reaction centres subunits of both photosystems, and an 8.5 kb operon encoding essential and non-essential genes. Finally, we tested degeneracy of the 51-codon genetic code by exploring the combinatorial design for the large subunit of RuBisCO, relying on restoration of photosynthesis in an *rbcL* mutant strain. More than 70 functional sequences with diverse codon adaptation indices were recovered. In all codon-compressed genes there was no observable penalty on photosynthetic growth.

**Significance Statement:** We demonstrate the use of a chloroplast genome as a synthetic biology platform to test modifications of the universal genetic code. Elimination of 13 of the 64 possible triplet codes (codons) was possible for several essential and/or highly expressed genes without affecting chloroplast function. The technology is mature, paving the way for codon compression of the entire genome and thus enabling further radical rearrangement or expansion of the genetic code.

## Introduction

The genetic code is universal and underpins all life on our planet. It provides the informational link between genes and their protein outputs, and therefore the instructions for building every organism. The code is simple and elegant - a set of 64 codons that code for 20 amino acids (AAs) and three stop commands - and, despite billions of years of evolution, has remained essentially unchanged among all extant life forms (1). The universality of the genetic code is perhaps surprising given that there are many ways in which AAs could be encoded using such a triplet code. Is the arrangement of the codon table the product of early optimisation during the evolution of protein translation, or is it a ‘frozen accident’, with other arrangements of the table equally functional? The assignment of codons is clearly non-random, with related residues typically occupying contiguous codon boxes in the table, and a simple analysis of the code shows it to be highly robust to mutational or translational errors. It is debatable whether this reflects a selective force or is just a consequence of the codon table’s early evolution, and indeed theoretical analyses have identified some of table variants more robust than the actual universal code (2, 3).

Testing completely different genetic codes is experimentally extremely challenging. However, improvements in custom DNA synthesis and DNA assembly in recent years have led to rapid advances in the field of synthetic genomics and allowed the design and synthesis of complete genomes for viruses, bacteria and yeast (4, 5). This has enabled moves towards modification of the genetic code. Successful examples of codon-compressed genomes have been reported, in which all instances of a particular codon have been modified to synonymous codons. These include removal of one and two of the stop codons in *E. coli* (6, 7), one stop codon in yeast Sc2.0 (8), and two sense and one stop codon in *E. coli* Syn61 (9). This enabled deletion of a previously essential tRNAs and release factors. Elimination of certain codons may then be used for reprogramming the unused codons, and indeed this has been done to incorporate non-canonical amino acids for biosynthesis of unnatural polymers (10). A further benefit came from resistance to viruses (10, 11) and improved biocontainment preventing horizontal gene transfer (12, 13).

Nonetheless, for the modified genomes assembled so far, even simple codon compression schemes proved to be difficult to implement. Out of eight schemes tested in *E. coli*, each introducing synonymous mutations in two sense codons to modify a 14.3 kb fragment encoding the cell division operon rich in essential genes, genes expressed at a range of levels and membrane proteins (14), only three were successful. When the successful scheme was applied to codon compression of the entire 3.9 Mb *E. coli* genome, which required modification of more than 18,000 codons, further troubleshooting and corrections were necessary to generate a viable strain (9). Design, assembly and testing of a 57-codon genome in *E. coli* (15) is still ongoing. Therefore, to assemble genomes with more radical codon compression and thus allow testing of reassignment and alternative genetic codes, much smaller and simpler genetic platforms are needed.

Chloroplast genomes are already highly reduced genetic systems, ranging between 120-200 kb depending on the species, and thus offer multiple advantages for testing codon compression schemes and reassignment. The 205 kb genome in the green alga *Chlamydomonas reinhardtii* encodes 70 protein genes (16), almost all of which have been annotated and characterized. High quality genome assemblies have been generated, and transcripts have been mapped (16, 17). Only three genes have introns and these have been successfully replaced with intron-less versions, and there are no overlapping genes, which would otherwise complicate recoding designs. Unlike in vascular plants, there is no RNA editing in the *C. reinhardtii* chloroplast (16) reducing the risk of affecting the editing process by modification of the coding sequences. The chloroplast genome can be engineered by homologous recombination allowing precise manipulation and constructs for engineering can be rapidly assembled by using Golden-Gate technology (18). Most significantly, *C. reinhardtii* can dispense with photosynthesis if supplied with acetate as a carbon source (19), meaning that radical alterations of its genome that affect function are not necessarily lethal.

The highly reduced nature of the chloroplast genetic system is also reflected in the small number of tRNA genes encoded by the genome and used for protein synthesis. Chloroplast genomes rely on wobbling and superwobbling of codon-anticodon interactions to read the entire codon table, where some tRNAs can recognise up to four codons. *In C. reinhardtii*, there are 29 tRNA genes, compared to more than 280 tRNA genes in the nucleus (20). In the absence of import of nucleus-encoded tRNAs into the chloroplast, the minimum number of tRNAs that is necessary to translate all 61 sense codons could be as low as 25 (two tRNAs for leucine, isoleucine, serine, arginine and methionine, and one for each of the remaining amino acids) (21). In this work we explored the potential to significantly compress the genetic code of the *C. reinhardtii* chloroplast genome. We took advantage of the simplicity of the chloroplast genetic system and designed one of the most radical codon compression schemes tested to date. We successfully replaced several key chloroplast coding sequences with the codon compressed variants and assessed the impact on chloroplast function.

## Results

### Establishing design principles for codon compression

Analysis of the codon usage of the *C. reinhardtii* chloroplast genome (**Fig. S1**) reveals a significant bias towards codons ending T or A, reflecting the high AT content (65.4%) of the plastome (16). Mapping the 29 tRNA genes (*Ct001-Ct029*) onto the codon table shows that the majority of amino acids are served by a single tRNA due the wobble and superwobble in codon recognition (21, 22). Two tRNA genes are duplicated (*trnA-UGC*, *Ct013* and *Ct023*, and *trnI-GAU*, *Ct012* and *Ct024*) due to their location in the 22 kb inverted repeat (IR). There are also two copies of *trnE-UU*C (*Ct003* and *Ct0016*), which are identical across the length of the tRNA gene, but the upstream and downstream flanking sequences are different. This might be due to the involvement of *trnE* in tetrapyrrole biosynthesis as well as protein synthesis, creating an increased demand for this tRNA (17, 23, 24). Among the non-duplicated tRNA genes reading codons from the same codon box, there are two tRNAs for methionine, one for initiation (*Ct026*) and one for elongation (*Ct018*), and two isoacceptor tRNAs for glycine, *trnG-GCC* (*Ct011*) and *trnG-UCC* (*Ct021*). Previous experiments in *Nicotiana tabacum* showed that *trnG-UCC* is essential and can read all GGN glycine codons by superwobbling, whereas *trnG-GCC* can be deleted (25). It should also be noted that the amber TAG stop codon is used in only 4 of the 70 protein coding genes and the opal TGA stop codon is not used at all.

The bias observed in the codon-tRNA landscape could be advantageous for compression of the genetic code as many of the rare codons could be replaced by frequently occurring synonymous codons without introducing a substantial number of codon changes in the genome. However, the reduced number of tRNAs in the chloroplast means that many of the rare codons are read by the same tRNA as other more frequently occurring codons within the same codon box. To allow reassignment of rare codons and avoid potential crosstalk, tRNAs could be modified to prevent wobbling. Alternatively, entire codon boxes could be modified, followed by deletion of the native tRNA gene.

We decided to take the latter approach for designing the genetic code compression scheme. For five amino acids (leucine, isoleucine, serine, arginine and glycine), there are two tRNAs with differing frequency of use. For example, *trnL-UAA Ct015*, which reads TTR codons (**Fig. 1**, marked in black), occurs 76% of the time, compared to 24% for *trnL-UAG Ct009* reading CTN codons (marked in grey). We hypothesized that for each of the five amino acids, the codons from the less frequently used codon box (target codons, marked in grey) could be changed to the most frequent codon(s) from the other box (marked in orange). After recoding all coding sequences in the chloroplast genome accordingly, together with changing the four amber (TAG) to ochre (TAA) stop codons, this would result in modification of 2136 codons (7.5% of all total codons) and the genetic code would be compressed to just 51 codons.

**Fig. 1.**
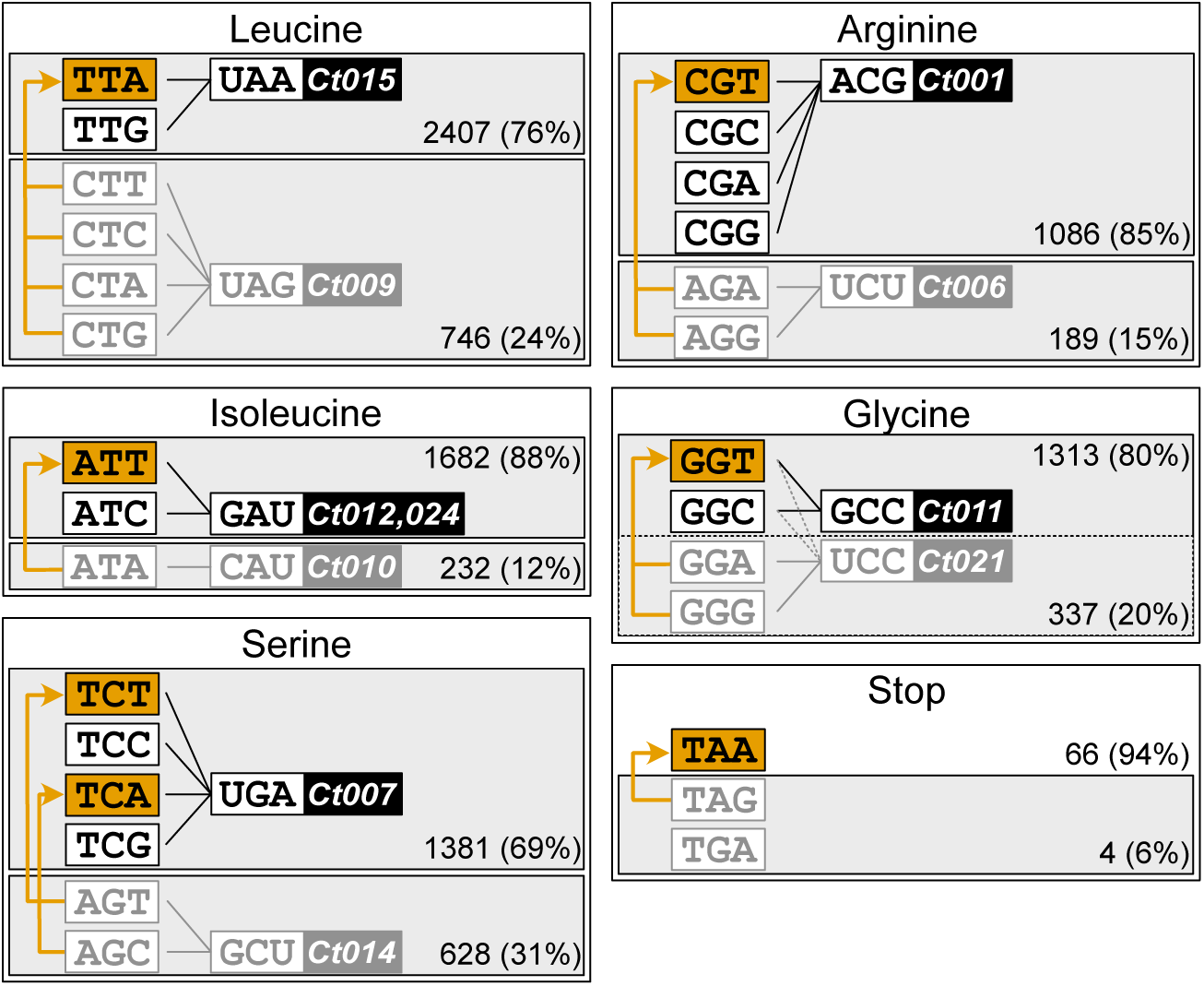
Proposed synonymous codon changes in the 51-codon defined compression scheme for the *C. reinhardtii* plastome. For each amino acid with two tRNAs with different anticodons (leucine, isoleucine, serine, arginine and glycine), the target codons from the less frequently used codon box (grey) are replaced by the most frequent codon(s) from the other codon box (orange). In addition, four *amber* stop codons (TAG) are replaced by the *ochre* codons (TAA); the third stop codon (*opal*, TGA) is not present in the *C. reinhardtii* plastome. Numbers indicate the occurrence and frequency of all codons from each codon box.

### Testing the 51-codon scheme by recoding essential genes

Most of the chloroplast-encoded protein genes are short, with a median length of 190 codons, and use on average only 40 out of 63 codons, meaning that many do not contain rare codons, and are therefore unsuitable proxies to infer the effects of the recoding schemes genome-wide (**Fig. S2A**). Of the larger genes, that of *rpoA* (738 codons) encoding the RNA polymerase subunit alpha, was chosen as the most suitable. It contains 57 unique codons, including at least one occurrence of 10 out of the 11 target sense codons for alternation in the compression schemes and has a frequency of these codons typical of the entire genome (7.5%, **Fig. S2B**). The gene is expressed as a monocistronic mRNA (16), so its manipulation would not affect other genes located upstream and downstream. Previously, a 345 bp fragment of the *rpoA* gene containing the promoter and 5’UTR was successfully replaced with the promoter and 5’UTR of *psbD* gene via homologous recombination, and at the same time the protein was internally tagged with the FLAG epitope sequence for immunodetection (26), showing that the gene can be easily manipulated. In *C. reinhardtii*, the plastid-encoded RNA polymerase (PEP) is the only RNA polymerase present in the chloroplast, as the nuclear genome does not encode any chloroplast-targeted RNA polymerase (NEP) (27, 28), therefore all *rpo* subunits are essential for both photosynthetic and heterotrophic growth (26). As such, the *rpoA* coding sequence (CDS) is a good target to test for the effect of recoding, since suboptimal or no expression of the gene would lead to decreased chloroplast gene transcription or no growth.

Three different codon-compressed *rpoA* coding sequences were designed and synthesised (**Fig. 2A; Supplementary Dataset 1**). The ‘defined compression’ scheme (cloned into plasmid pCSB246) replaced the 57 target codons, with the most frequent codons from the other codon boxes (marked in orange). To avoid a long 183 bp stretch of sequence homology between the native and the recoded gene that could act as a homology arm for recombination and potentially generate a chimeric WT-recoded gene, one other codon was modified (Lys387, rare AAG to frequent AAA, marked in green). The sequence included a FLAG-tag (blue box) to allow detection of the protein should homoplasmy not be reached (26). The other two sequence variants shown in **Fig. 2A** (pCSB247 and pCSB248) were designed as fully synthetic DNA sequences, generated using two different codon optimisation tools. ChimeraMAP (29) implemented in the CSO software (30) uses long substrings of codons that appear in the host coding sequences to retain genetic information embedded in the native sequence that conventional approaches cannot detect. Additionally, the Codon Usage Optimizer (CUO) software (https://github.com/khai-/CUO, accessed on 05-08-2020) generates the synthetic sequence based on codons and codon pairs frequencies of a handpicked set of highly expressed genes in the chloroplast of *C. reinhardtii*. Both synthetic sequences resulted in changes in 176 codons when compared to with the WT *rpoA* CDS, as some rare codons, both target (orange bars) and others (green bars), were preferentially replaced by more frequent codons. The sequences had a similar number of synonymous mutations (346 and 339, 15.4% and 15.1% of all bases respectively), but the distribution of the mutations was different, highlighting the contrasting approaches employed by the codon optimisation tools.

**Fig. 2.**
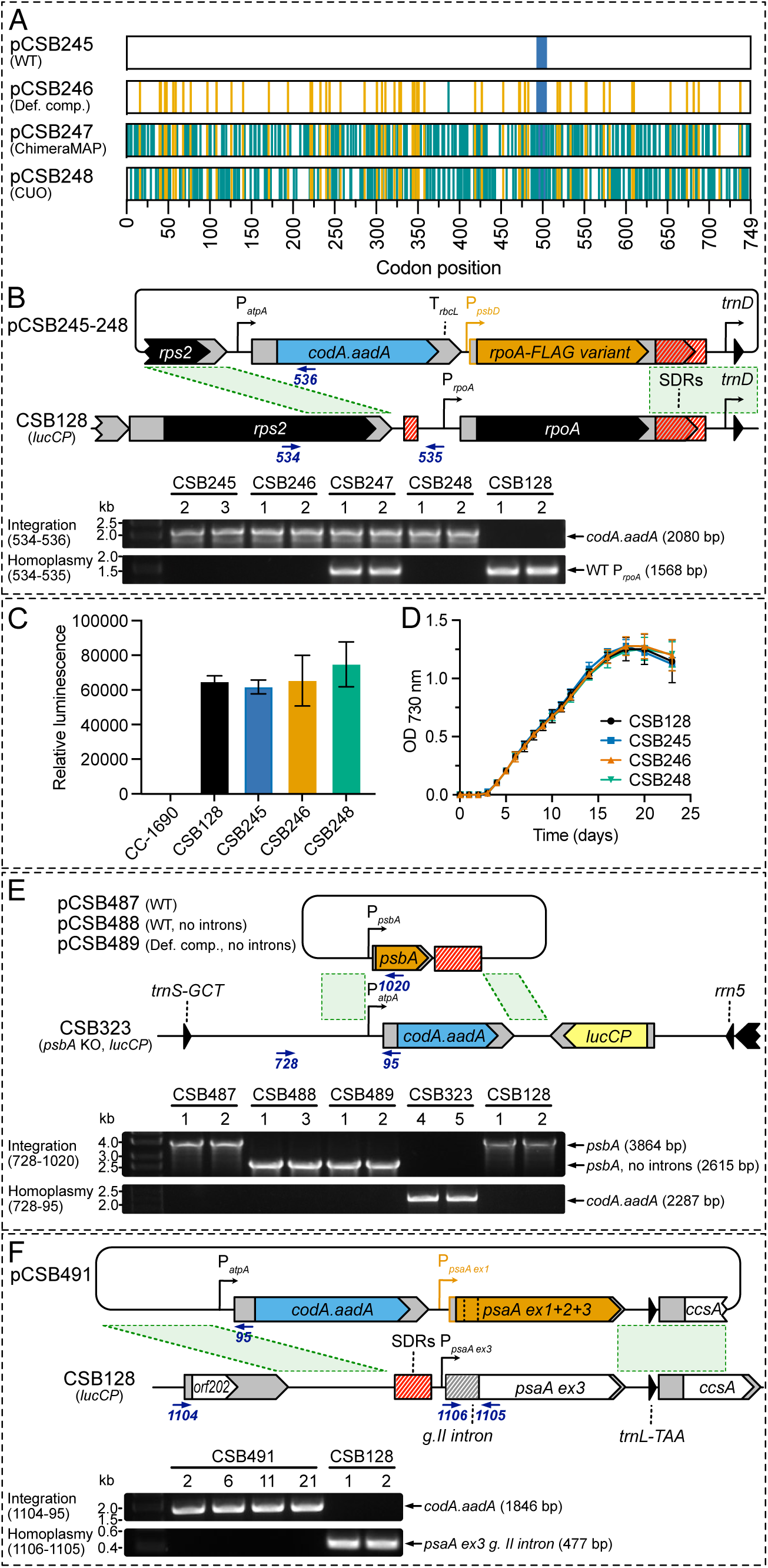
Codon compression of key genes in the *C. reinhardtii* plastome. **(A)** Schematic of *rpoA* sequences after codon compression by three different methods. Vertical orange lines represent synonymous mutations under the 51-codon defined compression scheme. Green lines represent other synonymous mutations (introduced to shorten the fragment of identity between the WT and defined compression variant or proposed by ChimeraMAP or CUO). The blue bar shows the position of the FLAG-tag. **(B) *Top:*** Schematic showing the replacement of the native *rpoA* coding sequence with the compressed variants encoded by plasmids pCSB246-248 via homologous recombination in the parental line CSB128 (which carries the *lucCP* cassette). Each plasmid replaced the 3.1 kb fragment between the left and right homology arms (green dashed lines) of the native *rpoA* gene and bordering Short Dispersed Repeats (SRDs, in red), with the *codA.aadA* cassette, and the recoded version of *rpoA* coding sequence, driven by the promoter/5’UTR of *psbD* gene. Also shown are the primers used for genotyping (blue arrows). ***Bottom:*** PCR of representative transformants using 534/536 primer pair to test for integration of the cassette or 534/535 primer pair to test for absence of parental band, *i.e.* homoplasmy. **(C)** Luciferase activity in homoplasmic transformants for CSB245, CSB246 and CSB248 together with CC-1690 and the parental line, CSB128. Cells were grown in TAP medium to mid-logarithmic phase, harvested, normalised to OD_730nm_ and activity measured using the Steady-Glo Assay System (Promega UK). **(D)** Phototrophic growth in cultures (25 mL) grown in HSM at light intensity of 80 μmol·m^-2^·s^-1^. **(E)** Strategy to express *psbA* with the 51-codon defined compression and without introns (CSB489). A native intron-containing sequence (CSB487) and a non-compressed control (CSB488) that just had the introns removed were also included. Plasmids pCSB487-489 were introduced into the *psbA* knock-out strain, CSB323, and transformants were selected by restoration of photosynthetic growth. **(F)** Strategy to express 51-codon compressed *psaA* gene. The three *trans*-spliced exons were combined, codon compressed and cloned under the *psaA ex1* promoter/5’UTR together with the *codA.aadA* cassette in plasmid pCSB491. After transformation into CSB128, the plasmid replaced the 3.1 kb fragment of the *psaA ex3* promoter, group II intron and coding sequence, and bordering SDRs.

These three codon-compressed *rpoA* sequences were used to generate a set of transformation plasmids (**Table S1**). Each plasmid included the *codA.aadA* cassette (31) upstream of the target gene and homology arms to replace the native *rpoA* CDS with the compressed variant (**Fig. 2B**). To avoid undesired recombination between the 919 bp fragment of *rpoA* promoter and 5’UTR (which is sufficiently long to act as a recombination homology arm, resulting in integration of the *codA.aadA* cassette without replacing the native *rpoA* CDS), the constructs used a 212 bp fragment of the *psbD* promoter and 5’UTR. A control plasmid pCSB245 had the non-recoded *rpoA* sequence but included the FLAG-tag. Plasmids were transformed into the chloroplast of *C. reinhardtii* strain CSB128 (**Table S2**) containing the *lucCP* firefly luciferase cassette integrated in the *rrn5-psbA* integration site in the inverted repeat (31). Spectinomycin-resistant transformants (16 to 22 independent lines per plasmid) were subcultured four times in increasing concentrations of antibiotics. Successful integration of the *codA.aadA* cassette was confirmed by colony PCR using a forward primer binding to the *rps2* gene located upstream of *rpoA* and a reverse primer binding to the *codA.aadA* cassette (**Table S3**), and homoplasmy was tested by the disappearance of the WT *rpoA* PCR product. Two representative transformants for each construct are presented in **Fig. 2B**, with the remaining shown in **Fig. S3**. The lines from the defined compression CSB246 constructs all successfully reached homoplasmy, whilst the frequency was 58% for the CUO CSB247 cell lines (**Table S4**). None were obtained for ChimeraMAP CSB248. Given the essential nature of the *rpoA* gene, it can be concluded that the first two designs allowed expression of functional enzymes, whereas ChimeraMAP did not.

The expression of the *lucCP* cassette integrated into the chloroplast genome of the parental strain CSB128 provided a proxy for the function of the chloroplast transcription-translation apparatus. CC-1690 (WT, parental strain of CSB128), CSB128 and homoplasmic lines of CSB245, CSB246 and CSB248 were grown in TAP medium, cells were harvested in the mid-logarithmic phase of growth and luciferase activity was tested (**Fig. 2C**). No statistically significant differences in luciferase activity were seen between the engineered strains. Moreover, photoautotrophic growth in minimal medium (HSM) was essentially identical between the homoplasmic lines and the CSB128 parental strain (**Fig. 2D**). This confirmed that recoding of *rpoA* in CSB246 and CSB248, as well as *rpoA* promoter/5’UTR replacement and incorporation of a FLAG-tag, did not affect the function of the RNA polymerase, chloroplast gene expression or cell growth under standard laboratory growth conditions.

One CSB246 line was re-transformed with plasmid pCSB322 (**Fig. S4A**) to restore the *rpoA* promoter and 5’UTR in front of the codon compressed *rpoA* CDS, using the *aphA6* antibiotic cassette for selection. Transformants were selected on agar plates supplemented with kanamycin and then re-streaked on medium supplemented with kanamycin and 5-fluorocytosine (to counterselect the *codA.aadA* cassette present in CSB246). Genotyping showed that cassette integration was homoplasmic in 15 out of 16 analysed CSB322 lines (**Fig. S4B** and **Table S4).** Complete recoding of *rpoA* was confirmed by next-generation amplicon sequencing (**Fig. S4C**) and a growth assay showed that the codon compressed *rpoA* CDS was also functional under the native promoter/5’UTR regulation (**Fig. S4D**).

The efficacy of the defined compression scheme was further confirmed by recoding of the *ycf1* gene, encoding a second essential protein, the Tic214 subunit of the translocon of the inner chloroplast membrane. This has 1996 codons and a frequency of 7.1% target codons (**Fig. S2B**). A total of 147 codons in the 6 kb *ycf1* gene were altered according to the 51-codon defined compression scheme with additional changes (**Fig. S5A, Supplementary Dataset 1**) and the synthesised DNA used to generate plasmid pCSB416 (**Fig. S5B**). Transformation of CSB128 with this construct was used to replace the WT *ycf1* CDS. Of 18 transformants selected for homoplasmy by restreaking, 14 were shown to be homoplasmic (**Fig. S5C** and **Table S4**), and sequencing confirmed replacement with the recoded gene (**Fig. S5D**). This indicated that the recoded gene was functional. There was no observable impact of the codon compression on phototrophic growth as assessed by measuring photoautotrophic growth in HSM (**Fig. S5E**).

### Recoding of highly expressed photosynthetic genes containing introns

We next investigated whether highly expressed genes could similarly be compressed, choosing those for the two reaction centre proteins of photosystems I and II. *psbA* encodes the D1 protein of PSII, which is the most highly expressed and rapidly turned-over protein in the chloroplast (32). The *psbA* gene is present in two copies in the genome (due to its location within the IR) and each has 4 introns, but these are not essential for *psbA* expression since the WT CDS can be replaced by an intron-less version (33). We aimed to replace the *psbA* CDS with an intron-less codon compressed variant containing 16 altered codons (4.5% of all codons). First, the non-photosynthetic *psbA* knockout strain CSB323 was generated by replacing the sole *psbA* gene with the *codA.aadA* cassette in the IR deletion mutant strain CSB358 (**Fig. S6**). As expected, the mutant was found to be non-photosynthetic. Then the mutation was complemented by transforming the knockout line with plasmids pCSB487-489 (**Fig. 2E**) encoding different variants of *psbA* (WT, WT with no introns and recoded with no introns, respectively) and plating on HSM minimal medium. Selection was by restoration of photosynthesis and transformants were obtained for all three plasmids, with the presence of the *psbA* CDS variant confirmed by PCR genotyping and sequencing. Homoplasmy was obtained for multiple transformants (**Fig. 2E, Fig. S7A**, **Table S4**).

PsaA, the A1 subunit of PSI, is also highly expressed, and is encoded by three *trans*-spliced exons distributed around the plastome, requiring an additional small chloroplast RNA *tscA* from a separate locus for splicing (34). The coding sequence of all three exons was combined and 44 target codons (5.9%) were modified using the defined compression scheme. Plasmid pCSB491 was designed and assembled to replace the *psaA* exon 3 group II intron and exon 3 coding sequence with the recoded variant of the full-length coding sequence (**Fig. 2F**). In the engineered lines CSB491 (**Fig. S8A)**, exons 1 and 2 are still present and transcribed, but they lack exon 3 group II intron for assembling the processed transcript, therefore the only functional transcript for translation of A1 protein is the recoded intron-less *psaA*. Growth assays in HSM minimal medium showed that homoplasmic lines encoding intron-less codon-compressed versions of both *psbA* and *psaA* were functional and recoding did not cause any defect in photosynthetic growth (**Fig. S7B and S8B**).

### Recoding of the trnM^e^-psbE-rps9-ycf4-ycf3-rps18-rps2 operon

Having successfully recoded single coding sequences without generation of chimeric WT-recoded products, we sought to codon compress a larger fragment of the chloroplast genome. As a test case we selected the 9.1 kb fragment located upstream of the already recoded *rpoA* gene, encoding a cluster of genes transcribed together from the *trnM^e^*promoter (**Fig. 3A**). Besides the essential *trnM*^e^, this operon also encodes multiple essential genes for plastid maintenance (*rps9*, *rps18* and *rps2*) and non-essential photosynthesis-related genes (*psbE*, *ycf4* and *ycf3*). Coding sequences of these six genes were codon-compressed according to the defined 51-codon scheme, with alterations to 94 target codons (**Supplementary Dataset 1**). To reduce regions of homology, up to 10 codons were altered at one or both ends of the coding sequences (**Fig. S9A**). The operon primary transcript and gene expression. Since mutations therein might result in altered expression of each gene, these sequences (up to 0.9 kb) were not modified so could result in generation of chimeric operons.

**Fig. 3.**
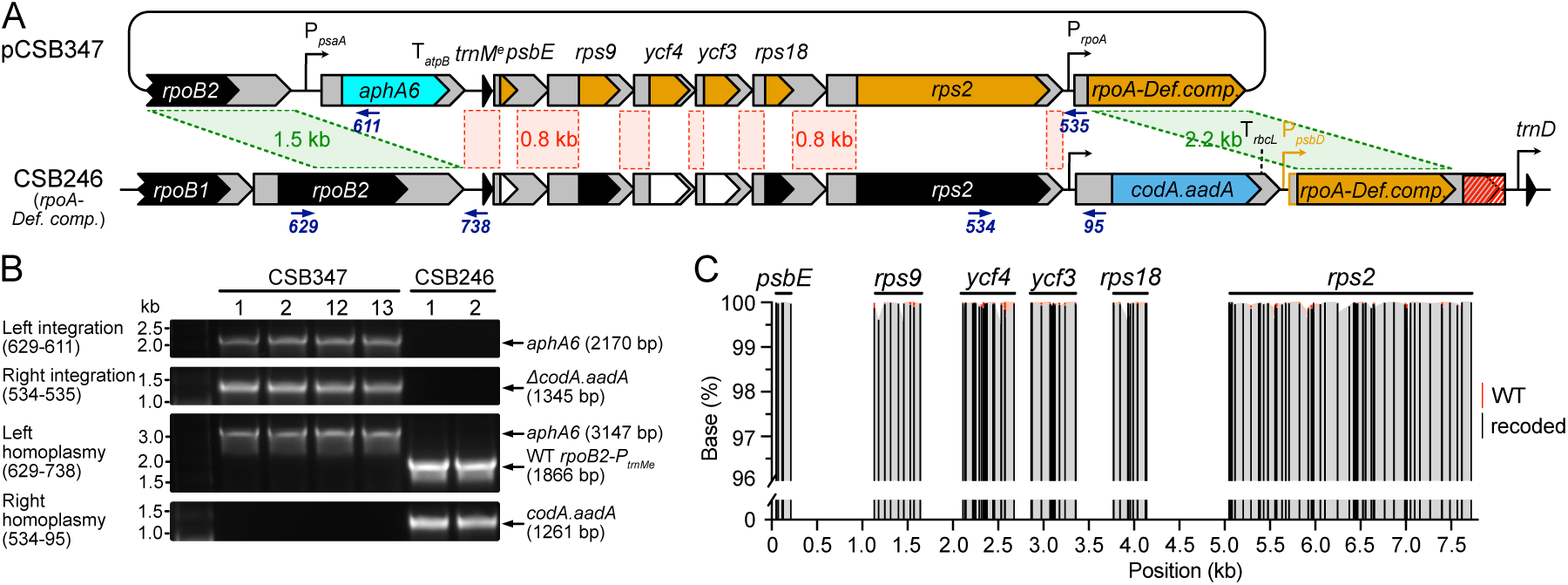
Codon compression of *trnM^e^-psbE-rps9-ycf4-ycf3-rps18-rps2* operon. **(A)** Schematic showing the process of recoding of the operon by transformation of the plasmid pCSB347 into the strain CSB246-1 (with *rpoA* compressed by the defined compression scheme). The plasmid replaced the 11.6 kb fragment between the 1.5 kb left and 2.2 kb right homology arms (green dashed lines) with the *aphA6* cassette and the recoded versions of the coding sequences. At the same time, the *codA.aadA* cassette was deleted and the *rpoA* promoter/5’UTR was restored upstream of the *rpoA* CDS. Red dashed lines represent non-coding fragments of the operon identical in pCSB347 and CSB246. Primers used for genotyping are represented by blue arrows. **(B)** PCR analysis of representative transformants confirming integration of the entire *trnM^e^* operon and homoplasmy. **(C)** Next Generation Sequencing of the *trnM^e^*operon in line CSB347-1. The operon sequence was PCR-amplified as sixteen 450 bp overlapping fragments with primers annealing outside of the modified codons and sequenced by Illumina Paired-End sequencing (Azenta, UK). Data represent the percentage of the recoded (black) and WT bases (red) at modified positions.

Plasmid pCSB347 encoding the recoded operon and the kanamycin resistance cassette was transformed into CSB246-1 (**Fig. 3A**). Homology arms in pCSB347 were designed in such a way that successful transformation would result in integration of the kanamycin cassette at the 5’ end of the operon and deletion of the *codA.aadA* cassette upstream of the already codon compressed *rpoA*. Therefore, transformants were selected on kanamycin and 5-fluorocytosine and restreaked twice to reach homoplasmy. 40 independent transformants were analysed and all showed correct integration of the kanamycin cassette, and 32 of them showed disappearance of the *codA.aadA* cassette (**Fig. 3B, Fig. S9B**, **Table S4**). Coding sequences of four representative lines were amplified and sequenced to confirm successful replacement of the WT sequences with the recoded versions. Additionally, one line was further analysed by deep next generation amplicon sequencing and no WT sequence was detected at the positions of target codons (**Fig. 3C**). Growth assay in HSM minimal medium showed that recoding of the operon did not cause any defects in photosynthetic growth under the conditions tested (**Fig. S9C**).

### Recoding of a highly expressed *rbcL* gene using a combinatorial codon approach

The recoding carried out so far changed the target codons to defined codons (indicated in orange in **Fig. 1**). However, even after compression to 51 codons, the genetic code remains degenerate, allowing translation from a single tRNA for each of serine and arginine from 4 different codons (TCN and CGN, respectively) and leucine, isoleucine and glycine from 2 different codons (TTR, ATY and GGY, respectively). In each case these have quite different codon usage frequencies, up to 90-fold (*e.g.* arginine CGT vs CGG; **Fig. S1**). We therefore sought to explore the effect of using each of the codon options in the compression scheme. We hypothesized that codon usage may be especially important for expression of highly abundant proteins, so chose the *rbcL* gene, encoding the large subunit of RuBisCO, as a test subject. This contains 26 redundant codons for four of the amino acids (22 leucine, 2 serine, 1 arginine and 1 glycine; **Fig. 4A**). Replacing these with all possible options from the 51-codon compression scheme (i.e. both orange and black codons in **Fig. 1**) would result in 5.4·10^8^ unique coding sequence variants. This would be virtually impossible to screen as independent cell lines, so instead, we decided to use a complementation assay to test a library of randomly generated codon-compressed variants of *rbcL* for their ability to rescue an *rbcL* mutant.

**Fig. 4.**
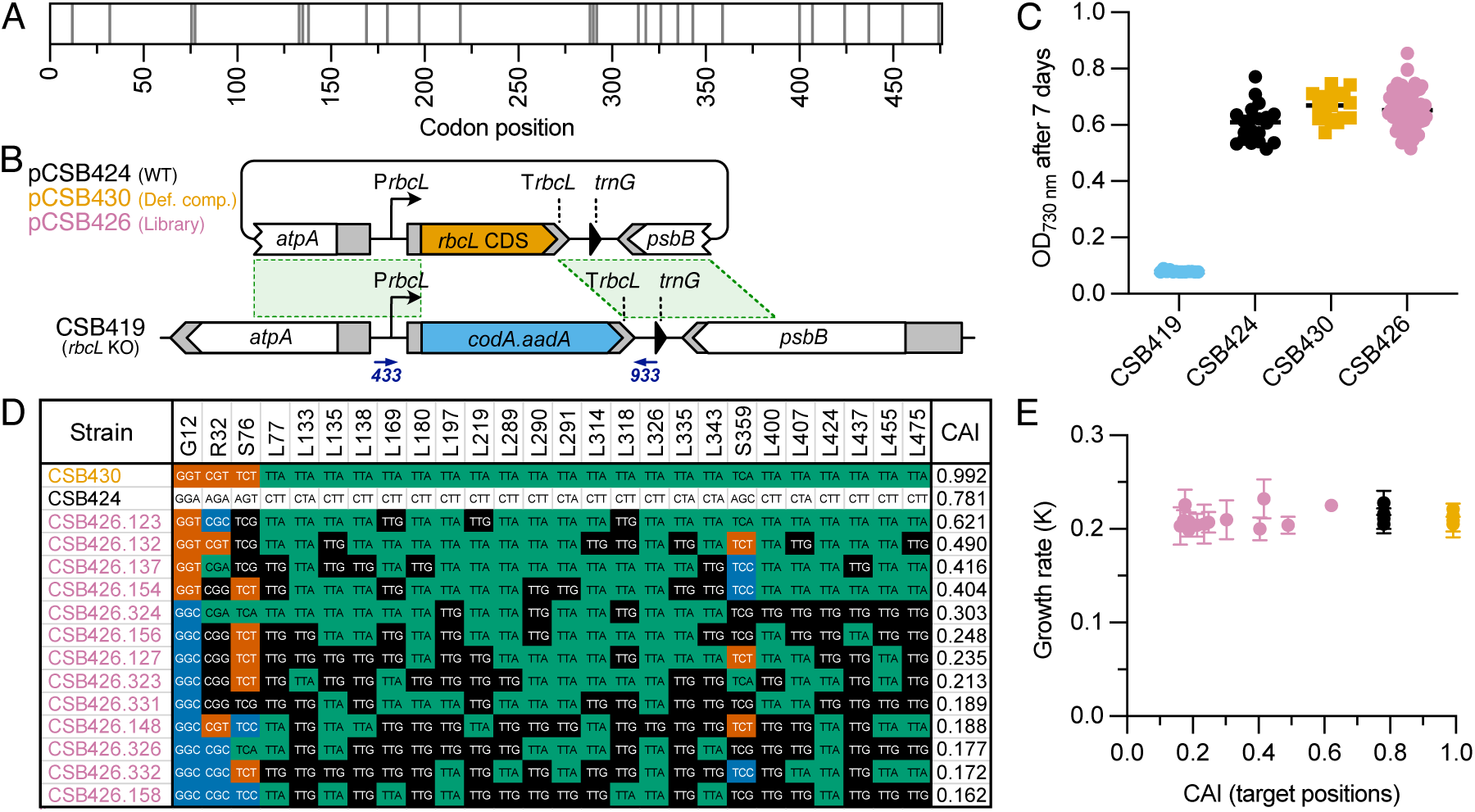
Codon compression of *rbcL* using the combinatorial approach. **(A)** Schematic of *rbcL* coding sequence containing 476 codons. Vertical grey lines represent positions of 26 target codons. **(B)** Schematic for recoding of *rbcL* coding sequence by complementation of the *rbcL* knockout strain CSB419 with plasmids pCSB424 (encoding the WT *rbcL* CDS), pCSB430 (defined compression scheme) or the pCSB426 plasmid library (combinatorially assembled *rbcL* variants). Primers used for genotyping and sequencing are represented by blue arrows. **(C)** Growth in minimal media of the *rbcL* KO (CSB419, n=13) and complemented strains (CSB424, n=20; CSB430, n=16 and CSB426, n=73). Cells were grown in 96 well plates (culture volume 200 μL) in HSM at light intensity of 80 μmol·m^-2^·s^-1^ and OD_730nm_ was measured after 7 days. Data points represent the average of three independent growth experiments. **(D)** Composition of the 26 target codons in 13 transformants selected from the CSB426 library compared to those in the WT *rbcL* (CSB424) and *rbcL* compressed using the defined compression scheme (CSB430). Sequences are sorted based on the decreasing value of the Codon Adaptation Index (CAI) calculated for the 26 target positions. Randomised codons are coloured by the third base of the codon: T - red, C - blue, A - green, G - black. **(E)** Growth rate of the CSB424 (n=4), CSB430 (n=4) and CSB426 (n=13) transformants in minimal medium plotted against the CAI of the target positions. Cultures (25 mL) were grown in flasks for 3 weeks. OD_730nm_ of cultures was measured every 24 h and the growth rate (K) was calculated after fitting data points to the Gompertz growth function (GraphPad Prism, v10). Error bars represent standard deviations for three independent cultures.

Firstly, an *rbcL* gene knockout line CSB419 (*rbcL* KO) was generated by found to be non-photosynthetic, and in this case light-sensitive. A library of complementation plasmids (pCSB426) was then generated by incorporation of synthetic degenerate oligonucleotides in which the *rbcL* sequence was randomized at the positions of the 26 target codons to introduce synonymous mutations (**Fig. S11**). Bulk sequencing of the plasmid library showed detectable degeneracy at the expected base positions (**Fig. S12**), with one exception, L326, where 99% were one codon (TTA). Control plasmids encoding the WT *rbcL* CDS (pCSB424) and the *rbcL* recoded using the defined compression scheme (pCSB430) were also assembled (**Fig. 4B**). These were transformed into CSB419 and lines encoding functional *rbcL* genes were selected by plating on minimal medium that supports only photosynthetic growth. Transformation efficiency was different between the control plasmids (40-80 colonies per µg DNA) and the pCSB426-lib (4-5 colonies per µg DNA), presumably reflecting the fact that not all library variants encoded a correctly assembled CDS. To reach homoplasmy, colonies were then re-streaked on minimal medium supplemented with 5-fluorocytosine, to counter-select the CSB419 chloroplast DNA containing the *codA.aadA* gene. After genotyping by PCR, the *rbcL* CDS was sequenced in 83 independent CSB426 lines. Of these, 10 had additional silent or missense substitutions at non-target positions and were eliminated from further analysis. The remaining 73 encoded unique *rbcL* sequences with synonymous mutations at the expected positions (**Fig. S13**). Photosynthetic growth, measured as OD_730nm_ after 7 days, was determined in a high-throughput experiment in 96-well plates in minimal media (**Fig 4C**). As expected, unlike the parental strain CSB419 (*rbcL* KO), the CSB424, CSB430 and CSB426 transformants were able to grow and showed similar end-point optical densities.

To characterize the complemented strains in more detail, the proportion of frequent (ending with A or T) to rare codons (ending with G or C) was examined They spanned the range of frequent:rare codon ratios from 21:5 to 8:18 and the set also included those showing the highest and the lowest OD_730nm_ values from the 96-well plate experiment and also sequences containing all possible permitted codons (four for serine and arginine and two for glycine and leucine) present at each randomized position. To measure the synonymous codon usage bias of selected sequences, the Codon Adaptation Index (CAI) (35) for the target positions was calculated (**Fig. 4D**). The selected lines were grown in 25 mL flask cultures for 3 weeks (**Fig. S13**) and the growth rates (K) for each line calculated and plotted against the CAI value (**Fig. 4E**). No significant differences between the growth rates of the codon compressed lines and the controls were observed, showing that compression of the genetic code for *rbcL* did not limit the photosynthetic growth of the recombinant strains.

## Discussion

We have taken the first step towards generation of the entire *Chlamydomonas* chloroplast genome with a radically compressed genetic code. We took advantage of the simplicity of the highly reduced genetic system of the plastome and designed the compression scheme to reduce the genetic code to just 51 codons. We tested the scheme by recoding and successfully replacing several native genes fulfilling different chloroplast functions, including essential genetic machinery genes (*rpoA, rps2, rps9, rps18*), an essential gene encoding a subunit of the membrane translocon complex (*ycf1*), and highly expressed genes encoding subunits of the photosynthetic complexes (*psaA, psbA, psbE* and *rbcL*). In total, we recoded more than 18 kb of coding sequences (21% of the plastome coding sequences) and modified 377 target codons (17.6% of all target codons in the plastome). Despite the significant compression of the genetic code of the key genes, none of the engineered strains suffered from loss of cell fitness, evidenced by the ease of reaching homoplasmy after transformation and WT-like growth under photoautotrophic conditions.

When designing the 51-codon compression scheme, we focused on targeting the codons for amino acids having two codon boxes and synonymously mutating the codons from the less frequently used codon box (2136 target codons, 7.5%). Using similar principles, even more radical codon compression schemes to just 46 codons are possible, (*e.g.* by mutating arginine CGN, isoleucine ATY and serine TCN codons) at a cost of larger number of codon swaps (5236 codons, 18.4%). Even further compression of the genetic code is suggested by the observation that several transgenes with compressed codons have been efficiently expressed in the *Chlamydomonas* chloroplast, including mVenus (31 unique codons), *Nluc* (29 codons) (36), and *lucCP* (31), which uses just 20 sense codons (one optimal codon for each amino acid). A codon compression scheme to just 20 sense codons has been previously proposed and used with moderate success for recoding of a small subset of essential yeast genes (22 out of 25 CDS remained genome, where gene families or genes with common domains would have repeated identical nucleotide sequences, remains unknown.

In codon compression schemes tested in *E. coli*, the target codons were replaced by codons with the closest match as calculated by the CAI, the tRNA adaptation index or translation efficiency (14), but using these metrics was not sufficient to predict optimal recoding schemes. In *Chlamydomonas*, the effect of using rare versus frequent codons on expression of the highly turned-over PsbA subunit has been investigated (39). Incorporation of certain rare codons (e.g. arginine CGG or AGG) impaired protein production, whereas the presence of others did not (e.g. glycine GGG or alanine GCG). In some cases (serine rare codon cluster 3xTCC), impairment of PsbA synthesis and ribosome pausing was observed, but only at high light intensities (1500 μmol photons·m^-2^·s^-1^), suggesting that translation speed is not a limiting factor under standard growth conditions (50-100 μmol photons·m^-2^·s^-1^). These effects are difficult to infer from the codon usage table and may also be context specific. Because of the highly biased chloroplast codon usage and significantly reduced set of tRNAs, where some tRNAs translate both frequent and rare codons due to superwobbling, our defined codon compression scheme replaced target codons with the most frequent permitted codon(s). Nevertheless, the discrepancy between the frequency of use between the target and recoded codons did not result in a growth phenotype in any of the recoded strains.

We explored this further using a combinatorial approach to identify which permitted codons are acceptable in the highly expressed photosynthetic gene, *rbcL*. After complementation of the *rbcL* knockout with the library of coding sequences, 73 unique functional sequences were identified (**Fig. 4C**). They contained all possible codons at all positions and showed different numbers of frequent and rare codons. However, none of the compressed sequences caused limitation of RbcL synthesis as shown by the growth assay. Also, mutating just 26 codons had only a minor effect on the CAI for the entire coding sequence (change from 0.77 to 0.70 in the variant encoding only rare codons). Because of the nature of the complementation experiment, only functional recoded sequences could be identified. In a situation when an attempt to recode a particular gene is unsuccessful, the native target codons could be included in the combinatorial design to identify positions that need troubleshooting.

The ease with which the 51-codon compression scheme could be introduced in the *Chlamydomonas* plastome is in striking contrast to studies testing codon compression schemes in *E. coli*. During the design of the *E. coli* Syn61 strain, additional identification and refactoring of 79 gene overlaps were necessary to allow their independent recoding (9). Some overlapping genes, such as *ftsI* and *murE*, were particularly difficult and required experimental troubleshooting by introduction of extended synthetic inserts to separate coding sequences and regulatory regions. Moreover, recoding of coding sequences of two hypothetical proteins (*yceQ* and *yaaY*) located within 5’UTRs of adjacent essential genes had negative effects on the regulation of these genes needing further redesign. In the end, the resulting strain had a 1.6-fold increase in the doubling time (9) indicating that recoding had reduced fitness overall. A different recoding project to generate an *E. coli* strain with a 57 codon genetic code required iterative genome synthesis and troubleshooting, with the resulting strain harbouring only maximum 39% of the recoded genome (40). It is likely that further compression of the genetic code in *E. coli* beyond 3-7 codons will be extremely technically challenging, if possible at all. In contrast, the small size, lack of overlapping genes and absence of cryptic sequences, as well as its non-essential nature, make the *Chlamydomonas* chloroplast genome a much more promising system for testing and implementation of radical codon compression schemes.

Recoding of *E. coli and* yeast genes showed that synonymous mutations in coding sequences may generate new sequences with promoter activities and induce intragenic transcription initiation resulting in generation of antisense transcripts (38, 40). These transcripts may affect gene regulation and active elimination of *de novo* promoters during genome design is necessary. Regulation of the gene expression in the chloroplast occurs not at the transcription initiation stage, but rather at the transcript processing stage, when different RNA binding proteins coordinate stabilisation, maturation and degradation of RNA (41, 42), possibly making the chloroplast gene regulation less sensitive to spurious promoters. Also, attempts to generate chloroplast synthetic promoters by randomisation of an existing strong promoter sequence gave only 10% of transformants showing detectable expression of the reporter gene (43). Generation of synthetic promoter libraries for AT-rich genomes showed that such organisms may have evolved more stringent recognition of promoter sequences to avoid initiation of transcription at spurious sites (44), and that would reduce a chance of generation of *de novo* promoters during codon compression of the entire genome. The effect of recoding on the chloroplast transcriptome will require further investigation in the future.

Beyond studies of the genetic code itself, there is considerable interest in generating a completely synthetic genome, and the small size of the plastome makes it attractive for this purpose. Previously, it was shown that chloroplast genomes can be assembled and propagated in heterologous hosts (45, 46). However, a successful replacement of the native genome with the synthetic version resulted in chimeric sequences likely due to the polyploidic nature of the chloroplast genome and the high level of homologous recombination (45). Efficient recombination can occur between fragments of identical sequence as short as 200 bp (31). We showed that recoding genes according to the defined 51-codon compression scheme makes the coding sequence sufficiently different to the native coding sequences and allows complete replacement of the genome fragments. In our initial experiments, we introduced additional mutations to reduce the lengths of identical sequences between the codon compressed and the native fragments (*e.g.* **Fig. 2A** and **Fig. S5A**). Analysis of a few randomly selected homoplasmic transformants did not reveal any chimeric sequences resulting from recombination. Subsequently, we found that even larger recoded fragments of ∼9 kb were efficiently incorporated, despite the presence of long unmodified non-coding fragments (**Fig. 3A**). Here, double selection was used by selecting for the integration of the positive marker (*aphA6*) at the left junction and deletion of a negative marker at the right junction (*codA*). This method could be used iteratively to replace the entire genome with recoded sequences, as in the assembly of the *E. coli* Syn61 genome (9, 14), although it would require development of a second negative marker for plastome engineering.

In conclusion, the simplicity of the chloroplast genome as a minimal genetic system offers a versatile synthetic biology platform for exploring compression, rearrangement and expansion of the genetic code. The results presented here put the chloroplast genome at the forefront of genetic code compression efforts and justify the synthesis of the entire recoded genome. Once the chloroplast genome is fully recoded, the five less frequently used tRNAs (*trnL-UAG Ct009, trnI-CAU Ct010, trnS-GCU Ct014, trnR-UCU Ct006 and trnG-UCC Ct021*) would become non-essential, so that they could be deleted. In the next step, the genetic code could be modified in different ways by reassigning the unused codons and functionally tested, addressing the question of whether the code is a product of early optimisation or the result of chance. Ultimately, the unused codons could also be reassigned to encode non-canonical amino acids for synthesis of new-to-nature proteins with novel properties.

## Materials and Methods

### Cultivation of *Chlamydomonas* strains

*Chlamydomonas reinhardtii* strains (**Table S2**) were grown mixotrophically or heterotrophically in TAP medium (47) or photoautotrophically in HSM (48), supplemented with trace elements (49), on plates with agar (2% w/v; Formedium Ltd, UK) or in liquid media, either at 25°C under constant illumination (80 μmol photons·m^-2^·s^-1^) or in darkness (photosynthetic mutant strains only). Cultures set up in cell culture flasks (Nunc EasYFlask 25 cm2, filter closure, Thermo Scientific, UK) were grown in shaking incubators at 120 rpm (Infors Multitron, Infors UK Ltd), while cultures in 96-well microplates (volume 200 μL) were grown in stationary incubators. Optical density of liquid cultures was measured using UV-VIS spectrophotometer (Thermo Scientific, UK) or CLARIOstar Plus plate reader (BMG Labtech, UK). Cell density was measured in C-Chip haemocytometer slides (Cambridge Bioscience, UK) using the Rebel hybrid microscope (Discover Echo, USA). Cultures were supplemented with spectinomycin (200-300 μg·mL^-1^; Melford, UK), kanamycin (200 mg·mL^-1^; Melford, UK) or 5-fluorocytosine (5 mg·mL^-1^; Thermo Scientific, UK) as detailed in the text.

### Plasmid design and assembly

Plasmids (**Table S1**) were generated using modular cloning methods. Recoded coding sequences were designed by manual replacement of the target codons or using codon optimization tools, ChimeraMAP (30) and CUO (Algal Biology and Biotechnology group, UCL; https://github.com/khai-/CUO, accessed on 05-08-2020). Synthetic DNA fragments were synthesised as gene fragments (Twist Biosciences, USA) or cloned ValueGENE plasmids (Azenta, UK). Constructs for transforming the chloroplast genome were assembled using the Start-Stop Assembly method (50) and included flanking regions for homologous recombination at the appropriate loci. Reactions mixtures were transformed into agar plates with respective antibiotics (carbenicillin, 100 μg·mL^-1^; tetracycline, 10 μg·mL^-1^). Plasmids were extracted and purified using Monarch® Miniprep Kit (NEB, UK) for small scale cultures, or GenEluteTM HP Midiprep Kit (Sigma-Aldrich, UK) for medium scale cultures. Plasmid concentrations were determined using a NanoDrop^TM^ spectrophotometer (Thermo Scientific, UK). All plasmids were verified by enzymatic digestion with appropriate restriction enzymes and sequence identity was confirmed by Sanger or Plasmid-EZ sequencing (Azenta, UK).

### Chloroplast transformation of *Chlamydomonas*

Transformation followed the protocol of (36) using bombardment with a Biolistics PDS-1000/HE device (Bio-Rad Laboratories, UK), with 1350 psi rupture discs (Bio-Rad Laboratories). Gold DNAdel carrier particles (Seashell Technology) were coated with the plasmid DNA at a concentration of 5 μg of DNA per 1 mg of gold particles, following manufacturer instructions. For each bombardment 2.5 μg of DNA was used. Transformants were selected on TAP agar plates supplemented with spectinomycin (200 μg·mL^-1^; Melford, UK) or kanamycin (200 μg·mL^-1^; Melford, UK), or on HSM agar plates (photosynthesis restoration). To reach homoplasmy, sixteen colonies per construct were streaked onto fresh selection plates, incubated for 7 days and then re-streaked up to 3 times. Genotypes of transformed strains were confirmed by using Phire Plant Direct PCR Kit (Thermo Scientific, UK) according to the manufacturer’s protocol. Transgene integration was verified using primers (**Table S3**) annealing to the transgene cassette and to the genome flanking the integration site. Homoplasmy was verified using primers that specifically amplify the wild-type gene at the integration locus, such that no product was formed once all chloroplast genome copies contain the transgene.

### Luciferase assay

Luciferase assays were performed using Steady-Glo Luciferase AssaySystem (Promega, UK). Samples of cultures in logarithmic phase of growth (1 mL) were centrifuged at 1000 x g for 10 minutes and pellets were resuspended in phosphate-buffered saline (PBS) to OD_730nm_ of 0.5. Samples (50 μL) were mixed with the lysis buffer (50 μL; 2 x Cell Culture Lysis Reagent Promega, UK, diluted in PBS) and incubated at room temperature for 10 minutes. Cell extracts (25 μL) were mixed with an equal volume of Steady-Glo® Reagent. Luminescence was measured at 580±80 nm using CLARIOstarTM plate reader after 5 min incubation.

### NGS Amplicon sequencing

Total DNA was extracted from cell pellets using the SDS method described by (51). Fragments of the chloroplast genome designated for NGS sequencing (450-500 bp) were amplified using the Platinum™ II Taq Hot-Start DNA Polymerase (ThermoFisher Scientific, UK) using primers with or without partial lllumina® adapter sequences (**Table S3**). PCR products were column purified using illustra™ GFX™ PCR DNA purification kit (VWR, UK). DNA concentration was determined using Qubit dsDNA HS Assay Kit (ThermoFisher Scientific, UK). Samples were normalised to 20 ng·μL^-1^, pooled and sequenced using Amplicon-EZ service (Azenta, UK). Raw reads were trimmed to quality score 20, mapped to the reference sequence using Unipro UGENE 52.0 BWA-MEM mapping tool and nucleotide variants were counted (52).

## Acknowledgments

We thank Dr. Katrin Geisler, Dr. Payam Mehrshahi, Dr. Gonzalo Mendoza-Ochoa, Dr. Lorraine Archer (University of Cambridge), and Dr. Henry Taunt (University College London) for helpful discussions.

## Author Contributions

P.M.M., H.O.J., S.P. and A.G.S. designed the research; P.M.M., K.C. and J.C. performed the research; P.M.M and A.H. analysed NGS sequencing data; P.M.M., H.O.J., S.P. and A.G.S. acquired the funding; P.M.M. and A.G.S. wrote the manuscript with input from all authors.

## Competing Interest Statement

The authors declare no competing interests.

## Notes

### Competing Interest Statement

The authors have declared no competing interest.

## References

1. E. V. Koonin, A. S. Novozhilov, Origin and evolution of the universal genetic code. Annu. Rev. Genet. 51, 45–62 (2017).

2. P. Błażej, M. Wnętrzak, D. Mackiewicz, P. Gagat, P. Mackiewicz, Many alternative and theoretical genetic codes are more robust to amino acid replacements than the standard genetic code. J. Theor. Biol. 464, 21–32 (2019).

3. Y. Omachi, N. Saito, C. Furusawa, Rare-Event Sampling Analysis Uncovers the Fitness Landscape of the Genetic Code. arXiv [q-bio.PE] (2022).

4. J. C. Venter, J. I. Glass, C. A. Hutchison 3rd, S. Vashee, Synthetic chromosomes, genomes, viruses, and cells. Cell 185, 2708–2724 (2022).

5. J. S. James, J. Dai, W. L. Chew, Y. Cai, The design and engineering of synthetic genomes. Nat. Rev. Genet. (2024). 10.1038/s41576-024-00786-y.

6. M. W. Grome, et al., Engineering a genomically recoded organism with one stop codon. Nature (2025). 10.1038/s41586-024-08501-x.

7. M. J. Lajoie, et al., Genomically recoded organisms expand biological functions. Science 342, 357–360 (2013).

8. Y. Zhao, et al., Debugging and consolidating multiple synthetic chromosomes reveals combinatorial genetic interactions. Cell (2023). 10.1016/j.cell.2023.09.025.

9. J. Fredens, et al., Total synthesis of Escherichia coli with a recoded genome. Nature 569, 514–518 (2019).

10. W. E. Robertson, et al., Sense codon reassignment enables viral resistance and encoded polymer synthesis. Science [Preprint] (2021). Available at: 10.1126/science.abg3029.

11. A. Nyerges, et al., A swapped genetic code prevents viral infections and gene transfer. Nature 615, 720–727 (2023).

12. R. E. B. Young, S. Purton, Codon reassignment to facilitate genetic engineering and biocontainment in the chloroplast of Chlamydomonas reinhardtii. Plant Biotechnol. J. 14, 1251–1260 (2016).

13. S. Changko, P. D. Rajakumar, R. E. B. Young, S. Purton, The phosphite oxidoreductase gene, ptxD as a bio-contained chloroplast marker and crop-protection tool for algal biotechnology using Chlamydomonas. Appl. Microbiol. Biotechnol. 104, 675–686 (2020).

14. K. Wang, et al., Defining synonymous codon compression schemes by genome recoding. Nature 539, 59–64 (2016).

15. N. Ostrov, et al., Design, synthesis, and testing toward a 57-codon genome. Science 353, 819–822 (2016).

16. S. D. Gallaher, et al., High-throughput sequencing of the chloroplast and mitochondrion of Chlamydomonas reinhardtii to generate improved de novo assemblies, analyze expression patterns and transcript speciation, and evaluate diversity among laboratory strains and wild isolates. Plant J. 93, 545–565 (2018).

17. M. Cavaiuolo, R. Kuras, F.-A. Wollman, Y. Choquet, O. Vallon, Small RNA profiling in Chlamydomonas: insights into chloroplast RNA metabolism. Nucleic Acids Res. 45, 10783–10799 (2017).

18. L. Esland, M. Larrea-Alvarez, S. Purton, Selectable Markers and Reporter Genes for Engineering the Chloroplast of Chlamydomonas reinhardtii. Biology 7 (2018).

19. R. Sager, S. Granick, Nutritional studies with Chlamydomonas reinhardi. Ann. N. Y. Acad. Sci. 56, 831–838 (1953).

20. V. Cognat, G. Pawlak, D. Pflieger, L. Drouard, PlantRNA 2.0: an updated database dedicated to tRNAs of photosynthetic eukaryotes. Plant J. 112, 1112–1119 (2022).

21. S. Alkatib, et al., The contributions of wobbling and superwobbling to the reading of the genetic code. PLoS Genet. 8, e1003076 (2012).

22. M. Fages-Lartaud, K. Hundvin, M. F. Hohmann-Marriott, Mechanisms governing codon usage bias and the implications for protein expression in the chloroplast of Chlamydomonas reinhardtii. Plant J. (2022). 10.1111/tpj.15970.

23. C. J. Howe, A. Smith, Plants without chlorophyll. Nature 349, 109–109 (1991).

24. A. C. Barbrook, C. J. Howe, S. Purton, Why are plastid genomes retained in non-photosynthetic organisms? Trends Plant Sci. 11, 101–108 (2006).

25. M. Rogalski, D. Karcher, R. Bock, Superwobbling facilitates translation with reduced tRNA sets. Nat. Struct. Mol. Biol. 15, 192–198 (2008).

26. S. Ramundo, M. Rahire, O. Schaad, J.-D. Rochaix, Repression of essential chloroplast genes reveals new signaling pathways and regulatory feedback loops in chlamydomonas. Plant Cell 25, 167–186 (2013).

27. S. S. Merchant, et al., The Chlamydomonas genome reveals the evolution of key animal and plant functions. Science 318, 245–250 (2007).

28. R. J. Craig, A. R. Hasan, R. W. Ness, P. D. Keightley, Comparative genomics of Chlamydomonas. Plant Cell 33, 1016–1041 (2021).

29. H. Zur, T. Tuller, Exploiting hidden information interleaved in the redundancy of the genetic code without prior knowledge. Bioinformatics 31, 1161–1168 (2015).

30. I. Weiner, Y. Feldman, N. Shahar, I. Yacoby, T. Tuller, CSO – A sequence optimization software for engineering chloroplast expression in Chlamydomonas reinhardtii. Algal Research [Preprint] (2020). Available at: 10.1016/j.algal.2019.101788.

31. H. O. Jackson, et al., CpPosNeg: A positive-negative selection strategy allowing multiple cycles of marker-free engineering of the Chlamydomonas plastome. Biotechnol. J. e2200088 (2022).

32. A. Lardans, et al., Biophysical, biochemical, and physiological characterization of Chlamydomonas reinhardtii mutants with amino acid substitutions at the Ala251 residue in the D1 protein that result in varying levels of photosynthetic competence. J. Biol. Chem. 273, 11082–11091 (1998).

33. U. Johanningmeier, S. Heiss, Construction of a Chlamydomonas reinhardtii mutant with an intronless psbA gene. Plant Mol. Biol. 22, 91–99 (1993).

34. U. Kück, O. Schmitt, The Chloroplast Trans-Splicing RNA-Protein Supercomplex from the Green Alga Chlamydomonas reinhardtii. Cells 10 (2021).

35. P. Puigbò, I. G. Bravo, S. Garcia-Vallve, CAIcal: a combined set of tools to assess codon usage adaptation. Biol. Direct 3, 38 (2008).

36. P. M. Mordaka, et al., Regulation of nucleus-encoded trans-acting factors allows orthogonal fine-tuning of multiple transgenes in the chloroplast of *Chlamydomonas reinhardtii*. Plant Biotechnol. J. (2024). 10.1111/pbi.14557.

37. Z. Liang, et al., Synthetic refactor of essential genes decodes functionally constrained sequences in yeast genome. iScience 25, 104982 (2022).

38. S. Jiang, et al., Building a eukaryotic chromosome arm by de novo design and synthesis. Nat. Commun. 14, 7886 (2023).

39. C. Weiss, I. Bertalan, U. Johanningmeier, Effects of rare codon clusters on the expression of a high-turnover chloroplast protein in Chlamydomonas reinhardtii. J. Biotechnol. 160, 105–111 (2012).

40. A. Nyerges, et al., Synthetic genomes unveil the effects of synonymous recoding. bioRxivorg 2024.06. 16.599206 (2024).

41. A. A. Zanini, M. F. Azim, T. N. McCray, T. M. Burch-Smith, “RNA binding proteins regulating chloroplast RNA metabolism” in *Nucleic Acids and Molecular Biology*, Nucleic acids and molecular biology., (Springer Nature Switzerland, 2024), pp. 39–74.

42. Y. Zhang, L. Tian, C. Lu, Chloroplast gene expression: Recent advances and perspectives. Plant Commun. 4, 100611 (2023).

43. R. Inckemann, et al., Advancing chloroplast synthetic biology through high-throughput plastome engineering of Chlamydomonas reinhardtii. bioRxiv 2024.05.08.593163 (2024).

44. P. M. Mordaka, J. T. Heap, Stringency of Synthetic Promoter Sequences in Clostridium Revealed and Circumvented by Tuning Promoter Library Mutation Rates. ACS Synth. Biol. (2018). 10.1021/acssynbio.7b00398.

45. B. M. O’Neill, et al., An exogenous chloroplast genome for complex sequence manipulation in algae. Nucleic Acids Res. 40, 2782–2792 (2012).

46. E. J. L. Walker, M. Pampuch, N. Chang, R. R. Cochrane, B. J. Karas, Design and assembly of the 117-kb Phaeodactylum tricornutum chloroplast genome. Plant Physiol. (2023). 10.1093/plphys/kiad670.

47. D. S. Gorman, R. P. Levine, Cytochrome f and plastocyanin: their sequence in the photosynthetic electron transport chain of Chlamydomonas reinhardi. Proc. Natl. Acad. Sci. U. S. A. 54, 1665–1669 (1965).

48. N. Sueoka, MITOTIC REPLICATION OF DEOXYRIBONUCLEIC ACID IN CHLAMYDOMONAS REINHARDI. Proc. Natl. Acad. Sci. U. S. A. 46, 83–91 (1960).

49. J. Kropat, et al., A revised mineral nutrient supplement increases biomass and growth rate in Chlamydomonas reinhardtii. Plant J. 66, 770–780 (2011).

50. G. M. Taylor, P. M. Mordaka, J. T. Heap, Start-Stop Assembly: a functionally Acids Res. (2018). 10.1093/nar/gky1182.

51. R. Zhang, et al., High-Throughput Genotyping of Green Algal Mutants Reveals Random Distribution of Mutagenic Insertion Sites and Endonucleolytic Cleavage of Transforming DNA. Plant Cell 26, 1398–1409 (2014).

52. K. Okonechnikov, O. Golosova, M. Fursov, UGENE team, Unipro UGENE: a unified bioinformatics toolkit. Bioinformatics 28, 1166–1167 (2012).

