## Supporting Information for "*Chlamydomonas* chloroplast genes tolerate compression of the genetic code to just 51 codons"

### Supporting Figures

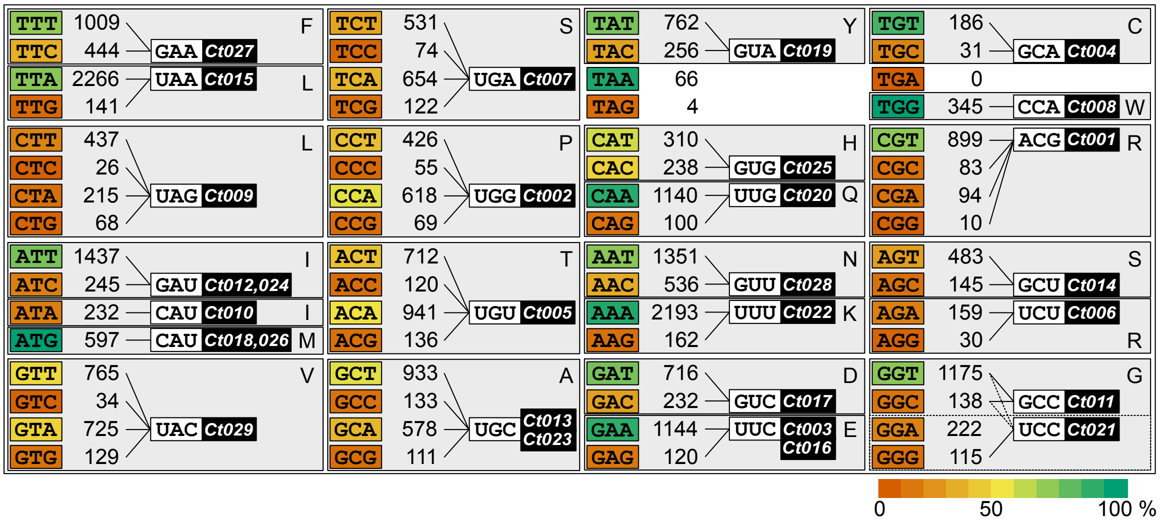


**Fig. S1. Codon table and tRNAs of *C. reinhardtii* chloroplast genome**. The absolute number of codons was calculated based on 70 unique open reading frames present in the chloroplast genome assembly (CPv4) for strain CC-503 (1). tRNA genes and anticodons were predicted by tRNAscan-SE 2.0 (2) and cross-referenced with the PlantRNA database (3). tRNA gene identifiers (*Ct001-029*) were taken from Cpv4 (1). Codon-anticodon interactions were predicted based on wobbling and superwobbling rules established experimentally for the plastome of *Nicotiana tabacum* (4) and bioinformatics analysis for *C. reinhardtii* (5). The heat-map represents the percentage of each codon among all codons encoding the same amino acid.

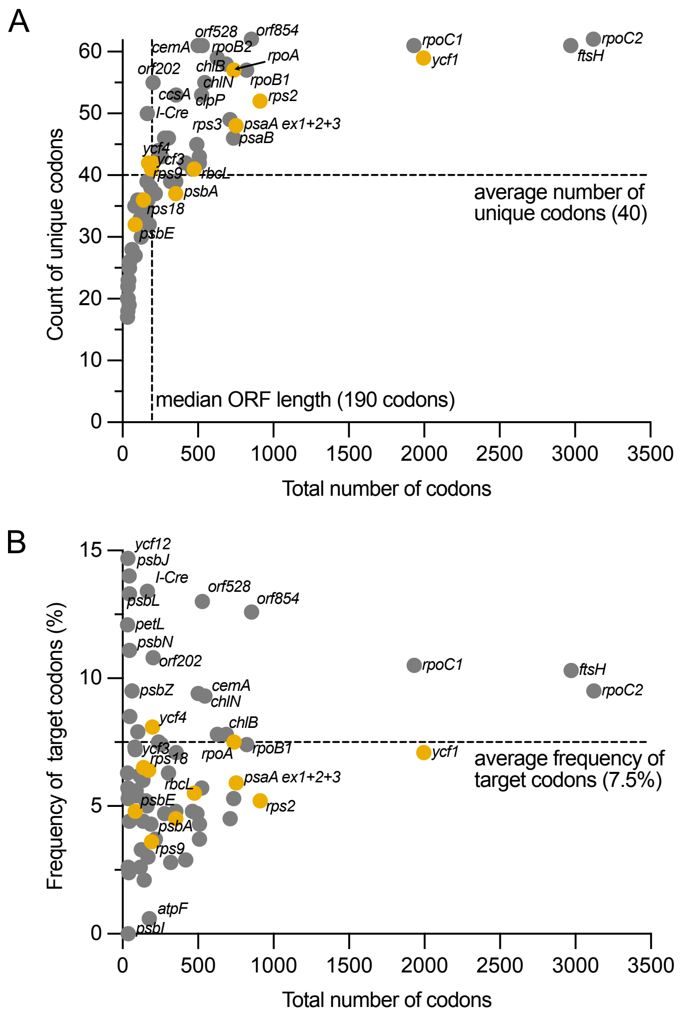


**Fig. S2.** Count of unique codons (**A**) and frequency of 51-codon compression scheme target codons (**B**) against the total number of codons in protein coding sequences in the *C. reinhardtii* chloroplast genome. Genes successfully recoded in this study are marked in orange.

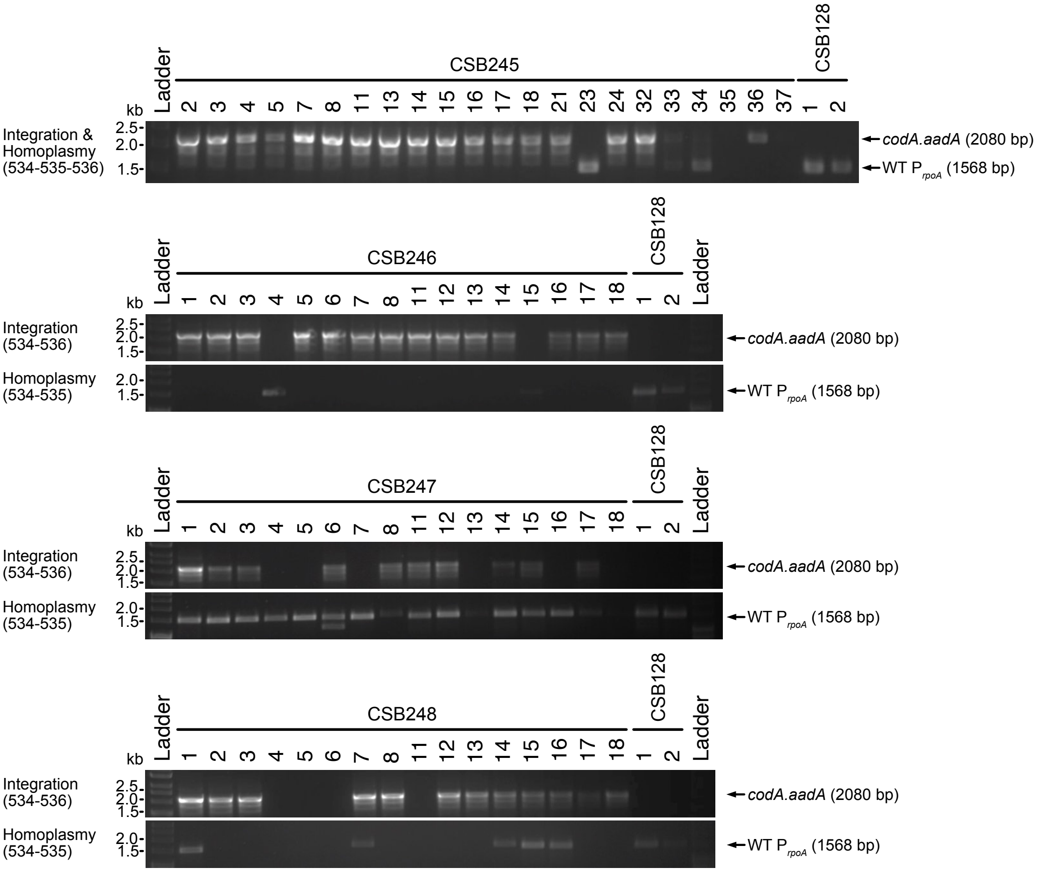


**Fig. S3. Genotyping of *rpoA* recoding transformants.** Integration of the *codA.aadA* cassette and homoplasmy was tested by PCR for lines transformed with plasmids pCSB245, pCSB246, pCSB247 and pCSB248. Primers used for amplification are listed on the left and sizes of the expected products on right. Parental strain CSB128 was used as a control. Ladder - HyperLadderTM 1kb (Bioline).

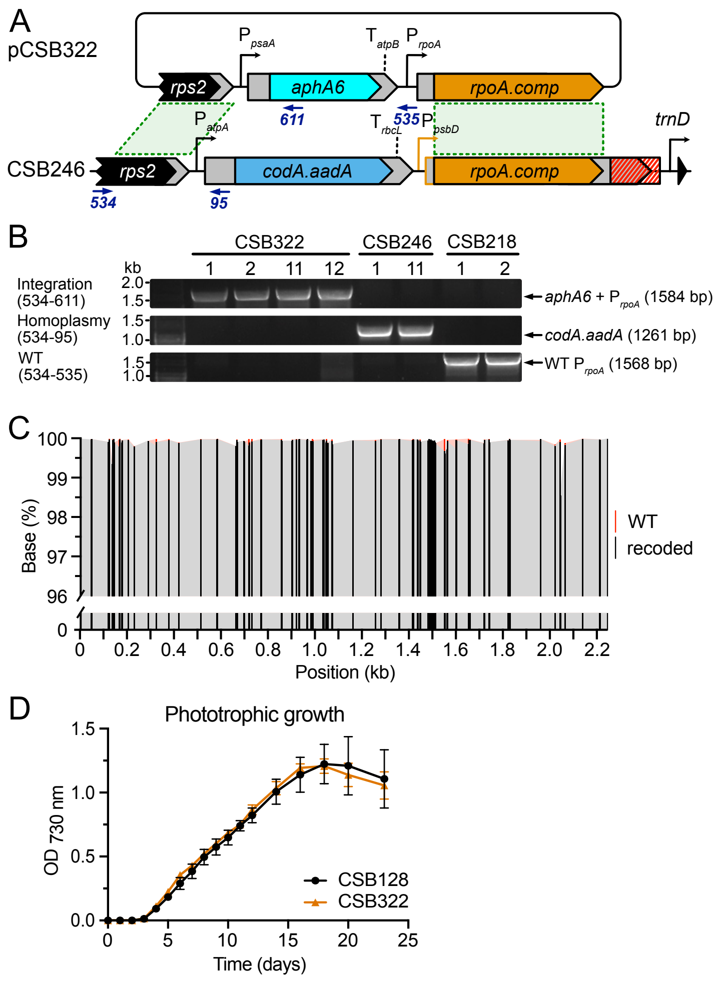


**Fig. S4. Restoration of *rpoA* promoter and 5'UTR in CSB246. (A)** Schematic showing the restoration of the *rpoA* promoter and 5'UTR upstream of the *rpoA* CDS recoded using the 51 defined compression scheme. The *rpoA* codon-compressed strain CSB246 was transformed with plasmid pCSB322 and transformants were selected on kanamycin and 5-FC. Primers used for genotyping are represented by blue arrows. **(B)** Genotyping of the representative transformant lines. Primers oligoCSB534 and oligoCSB611 were used for confirmation of the plasmid integration resulting in a product of 1584 bp. Primer pairs oligoCSB534-oligoCSB95 and oligoCSB534-oligoCSB535 were used for detection of the CSB246 and WT plastome DNA (products of 1261 bp and 1568 bp respectively). **(C)** Confirmation of *rpoA* recoding by Amplicon NGS. The entire coding sequence of *rpoA* was amplified as five overlapping 497 bp PCR products, pooled and pair-end sequenced using Illumina sequencing platform (Amplicon-EZ, Azenta). Bars represent positions of synonymous substitutions. Frequency of bases: black - recoded, red - WT. **(D)** Phototrophic growth of homoplasmic CSB322 lines in cultures (20 mL) grown in HSM at light intensity of 80 µmol·$m^{-2}$·$s^{-1}$.

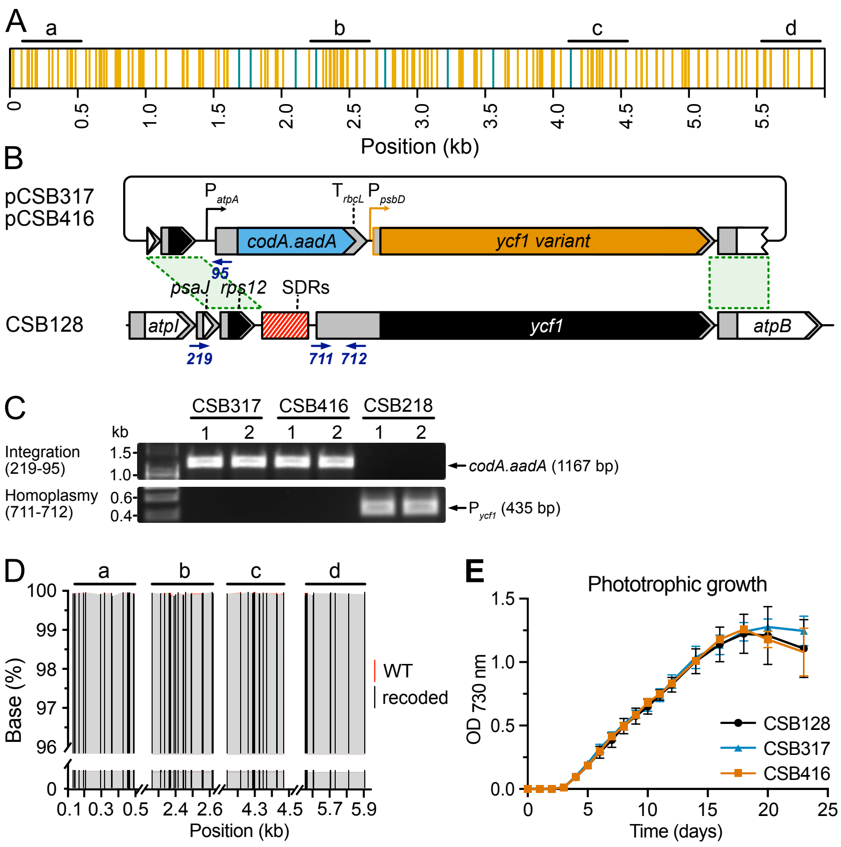


**Fig. S5. Codon compression of the *ycf1* coding sequence. (A)** Schematic showing the positions of target codons in the *ycf1* coding sequence. Vertical lines represent synonymous mutations in the coding sequence (orange - target codons, green - other codons). (**B)** Plastome recombination with plasmid pCSB416 used for recoding of the *ycf1* gene coding sequence. The plasmid replaced the 8148 bp fragment of the plastome located between the left and right homology arms (green dashed lines) containing Short Dispersed Repeats (SDRs, in red), promoter, 5'UTR and coding sequence of *ycf1* gene with the codA.aadA cassette, promoter and 5'UTR of *psbD* gene, and the recoded version of *ycf1* coding sequence. Plasmid pCSB317 was used as control, where the SDRs and ycf1 promoter/5'UTR were replaced with the codA.aadA cassette and the promoter and 5'UTR of *psbD* gene. **(C)** Genotyping of the transformant lines. Primers oligoCSB219 and oligoCSB95 were used for confirmation of the plasmid integration resulting in a product of 1167 bp. Primers oligoCSB711 and oligoCSB712 were used for detection of the WT DNA (product of 435 bp). **(D)** Confirmation of *ycf1* recoding by Amplicon Next Generation Sequencing. Four fragments of the *ycf1* coding sequence were amplified as 450 bp PCR products, pooled and pair-end sequenced using Illumina sequencing platform (Amplicon-EZ, Azenta). Bars represent silent substitutions. Frequency of bases: black - recoded, red - WT. **(E)** Phototrophic growth of the homoplasmic transformant lines. CSB128 and transformants CSB317 and CSB416 were grown in HSM at light intensity of 80 µmol·m$^{-2}$·s$^{-1}$.

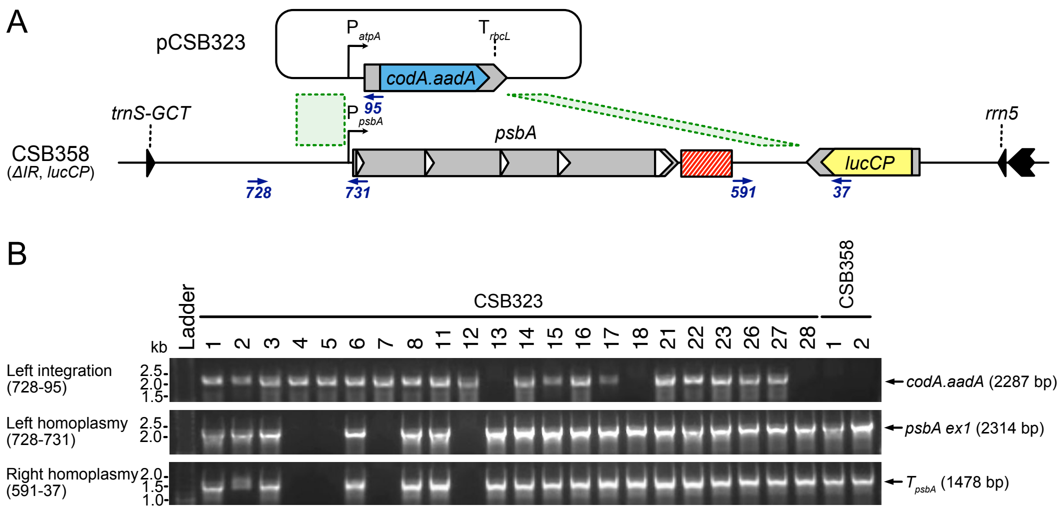


**Fig. S6. Generation of the *psbA* knockout strain. (A)** Schematic showing the process of deletion of the *psbA* gene by transformation of the plasmid pCSB323 into the strain CSB358 (IR deletion strain in CSB128 *lucCP*). The plasmid replaced the 7.8 kb fragment between the left and right homology arms (green dashed lines) with the *codA.aadA* cassette. Primers used for genotyping are represented by blue arrows. **(B)** PCR analysis of transformants confirming homoplasmic deletion of *psbA*. Primers oligoCSB728-oligoCSB95 were used for confirmation of the plasmid integration resulting in a product of 2287 bp. Primer pairs oligoCSB728-oligoCSB731 and oligoCSB591-oligoCSB37 were used for detection of the WT DNA on left and right junctions (products of 2314 bp and 1478 bp respectively).

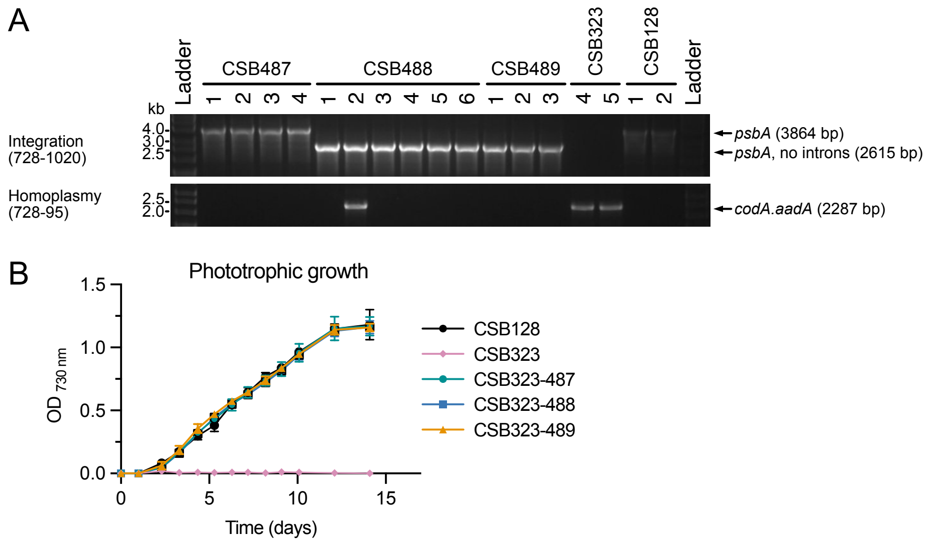


**Fig. S7. Codon compression of *psbA*. (A)** PCR analysis of transformants confirming reinsertion of the modified *psbA* in CSB323. Primers oligoCSB728 and oligoCSB1020 were used for detection of the *psbA* coding sequence resulting in products of 3864 bp (with introns) and 2615 bp (without introns). Primers oligoCSB728 and oligoCSB95 were used for homoplasmy screening by detection of the *codA.aadA* cassette (product of 2287 bp). **(B)** Phototrophic growth of the homoplasmic complemented lines and the parental lines. Cultures (n=3) were grown in HSM at light intensity of 80 µmol·$m^{-2}$·$s^{-1}$.

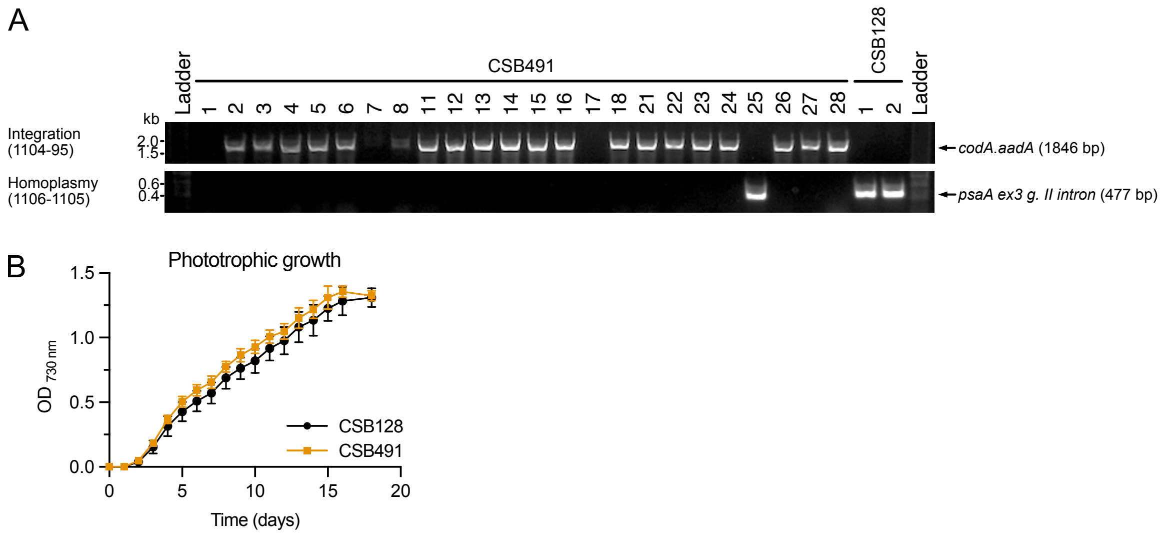


**Fig. S8. Codon compression of *psaA*. (A)** PCR analysis of transformants confirming recoding of *psaA* in CSB128. Primers oligoCSB1104 and oligoCSB95 were used for integration of the *codA.aadA* cassette resulting in a product of 1846 bp. Primers oligoCSB1106 and oligoCSB1105 were used for homoplasmy screening by detection of the *psaA ex. 3* group II intron (product of 477 bp). **(B)** Phototrophic growth of the homoplasmic complemented lines and the parental lines. Cultures (n=8) were grown in HSM at light intensity of 80 µmol·m$^{-2}$·s$^{-1}$.

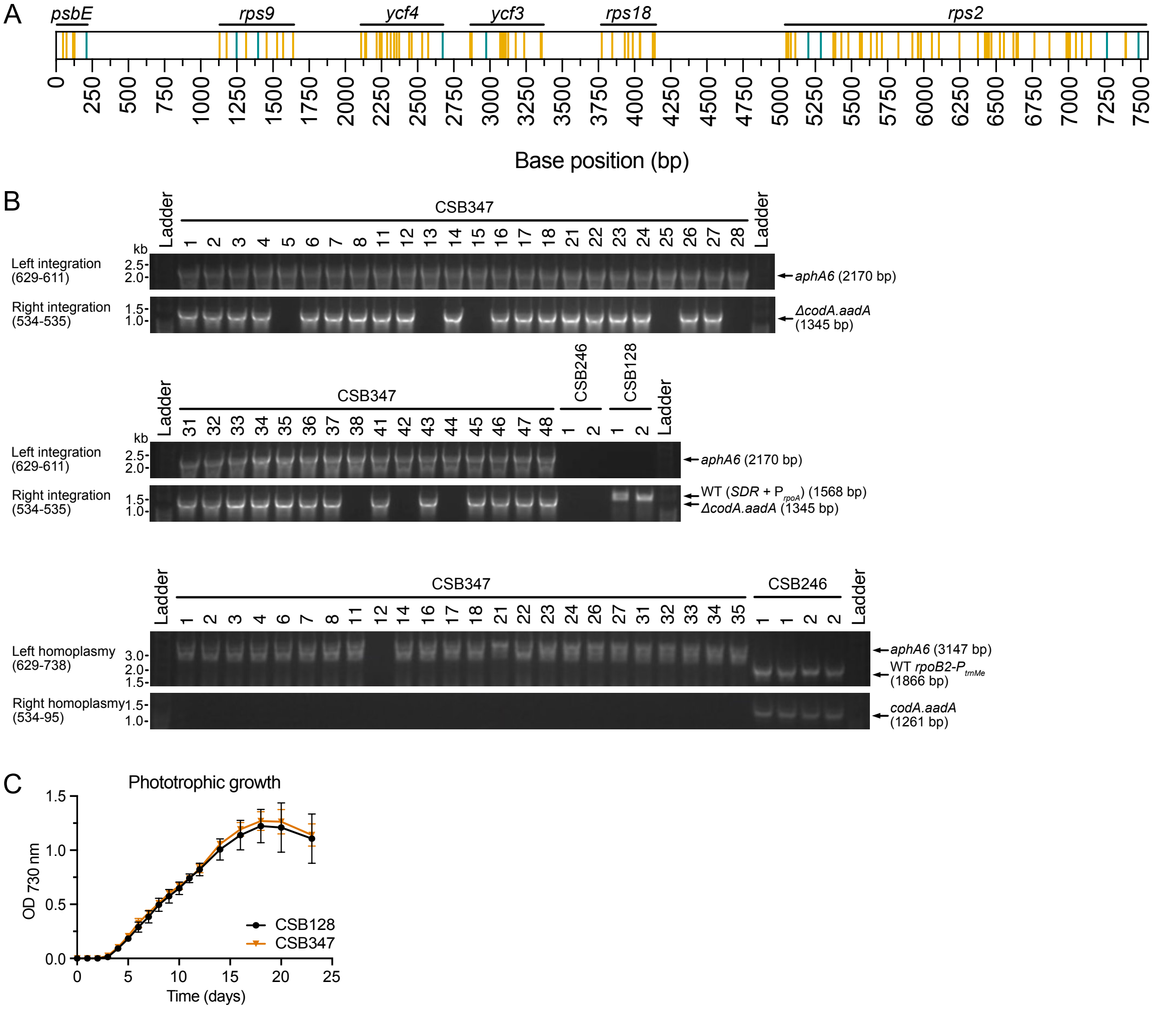


**Fig. S9. Codon compression of the *trnM^e^-psbE-rps9-ycf4-ycf3-rps18-rps2* operon (A)** Schematic showing the positions of target codons in the *trnM^e^* operon. Coding sequences of individual genes are marked above. Vertical lines represent synonymous mutations in the coding sequences (orange - target codons, green - other codons). **(B)** PCR analysis of transformants confirming integration of the entire *trnM^e^* operon (oligoCSB629-oligoCSB611 for the left integration junction and oligoCSB534-oligoCSB535 for the right integration junction) and homoplasmy screen (primers oligoCSB629-oligoCSB738 for the left junction homoplasmy and oligoCSB534-oligoCSB95 for the right junction homoplasmy). **(C)** Phototrophic growth of the representative homoplasmic lines and the parental line CSB128. Cultures (n=4) were grown in HSM at light intensity of 80 $\mu mol \cdot m^{-2} \cdot s^{-1}$.

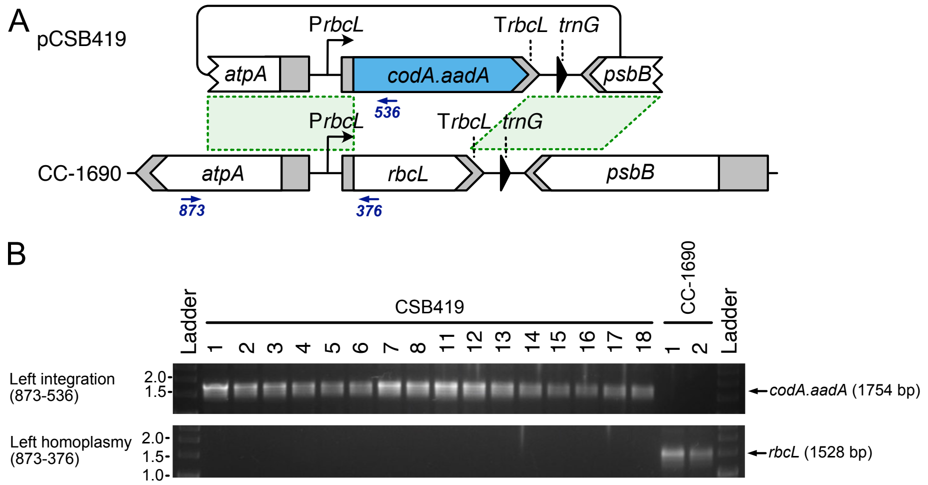


**Fig. S10. Generation of the *rbcL* knockout strain. (A)** Schematic showing the process of deletion of the *rbcL* gene by transformation of the plasmid pCSB419 into the strain CC-1690. The plasmid replaced a 1.4 kb fragment between the left and right homology arms (green dashed lines) with the *codA.aadA* cassette. Primers used for genotyping are represented by blue arrows. **(B)** PCR analysis of transformants confirming homoplasmic deletion of *rbcL*. Primers oligoCSB873 and oligoCSB536 were used for confirmation of the plasmid integration resulting in a product of 1754 bp. Primers oligoCSB873 and oligoCSB376 were used for the homoplasmy screen by detection of the *rbcL* CDS (product of 1528 bp).

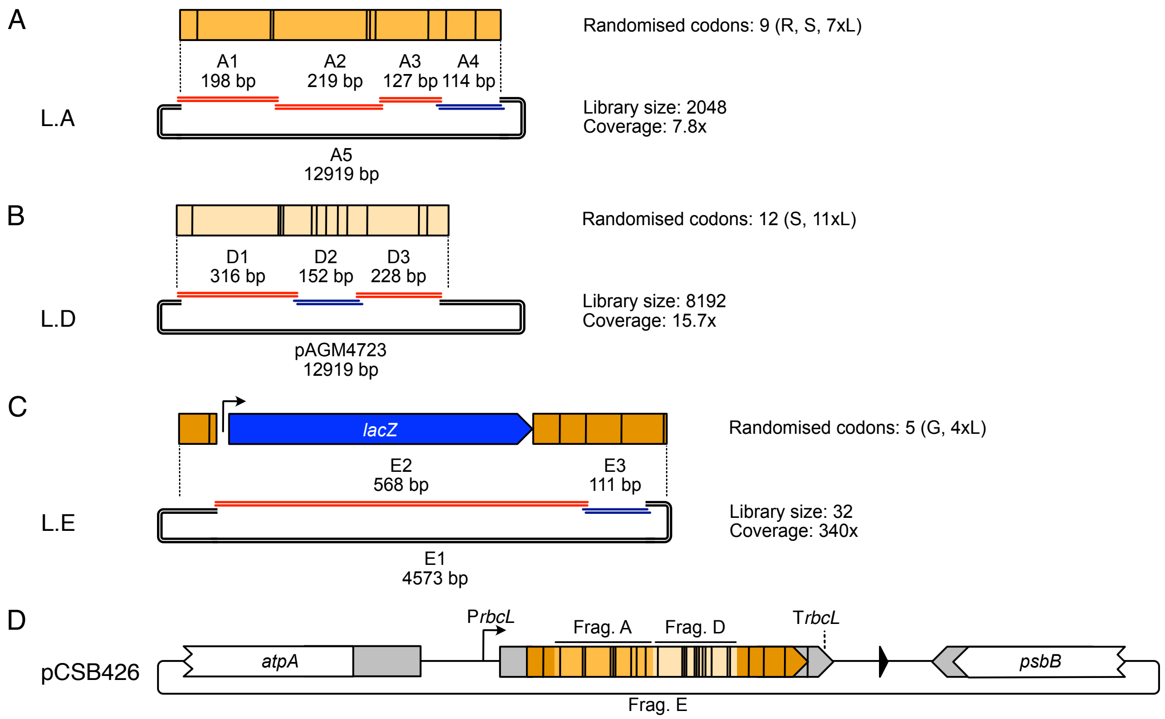


**Fig. S11. Generation of the *rbcL* combinatorial library pCSB426.** Plasmid library pCSB426 was constructed from three libraries of smaller sizes, L.A **(A)**, L.D **(B)** and L.E **(C)**, each encoding a different fragment of the *rbcL* coding sequence. The smaller libraries were generated by amplification of DNA fragments with degenerate nucleotides in the primer overhangs (fragments marked in red) or by annealing of degenerate single strand oligonucleotides (fragments marked in blue). Oligonucleotides used for amplification and annealing are in Table S3. Black vertical lines represent positions of the degenerate bases. Fragments were mixed, ligated by Golden-Gate reactions with BsmBI (L.A) or BbsI (L.D and L.E) and transformed to electrocompetent NEB® 10-beta *E. coli* cells. Number of randomised codons, the size of each library and the obtained coverage are indicated on the right side of each schematic. **(D)** The pCSB426 library was generated by mixing libraries L.A, L.D and L.E, ligation by Golden-Gate reaction with BsaI and transformation to electrocompetent NEB® 10-beta *E. coli* cells. Transformants were inoculated into 100 mL of LB, incubated overnight at 37°C and plasmid DNA was extracted using GenEluteTM HP Midiprep Kit (Sigma-Aldrich, UK).

| 3rd base | G12 | R32 | S76 | L77 | L133 | L135 | L138 | L169 | L180 | L197 | L219 | L289 | L290 | L291 | L314 | L318 | L326 | L335 | L343 | S359 | L400 | L407 | L424 | L437 | L445 | L475 |
|---|---|---|---|---|---|---|---|---|---|---|---|---|---|---|---|---|---|---|---|---|---|---|---|---|---|---|
| A | 0 | 20 | 20 | 38 | 57 | 44 | 30 | 55 | 50 | 44 | 25 | 60 | 62 | 70 | 44 | 43 | 99 | 45 | 37 | 8 | 69 | 79 | 51 | 36 | 39 | 25 |
| G | 0 | 50 | 31 | 62 | 43 | 56 | 70 | 44 | 50 | 56 | 75 | 34 | 30 | 29 | 56 | 57 | 0 | 55 | 63 | 30 | 31 | 21 | 49 | 64 | 61 | 75 |
| T | 34 | 16 | 26 | 0 | 0 | 0 | 0 | 1 | 0 | 0 | 0 | 6 | 9 | 2 | 0 | 0 | 0 | 0 | 0 | 30 | 0 | 0 | 0 | 0 | 0 | 0 |
| C | 66 | 14 | 23 | 0 | 0 | 0 | 0 | 0 | 0 | 0 | 0 | 0 | 0 | 0 | 0 | 0 | 1 | 0 | 0 | 33 | 0 | 0 | 0 | 0 | 0 | 0 |

**Fig. S12. Confirmation of degeneracy at the target codons positions in pCSB426 library.** Sanger sequencing chromatograms of *E. coli* purified pCSB426 libraries were analysed using EditR 1.0.10 (6) and frequency of base at 3rd position of each target codons was calculated. The expected values were 25% of A, G, T and C for arginine and serine, 50% of T and C for glycine and 50% of A and G for leucine. Analysis showed significant degeneracy at target codon positions except for leucine 326, which was A in 99% of codons. The bases in 1st and 2nd positions did not show any degeneracy (data no shown). Data represent average frequencies for three independently assembled and sequenced pCSB426 libraries.

| | G12 | R32 | S76 | L77 | L133 | L135 | L138 | L169 | L180 | L197 | L219 | L289 | L290 | L291 | L314 | L318 | L326 | L335 | L343 | S359 | L400 | L407 | L424 | L437 | L455 | L475 | 3rd base A or T | 3rd base G or C |
|---|---|---|---|---|---|---|---|---|---|---|---|---|---|---|---|---|---|---|---|---|---|---|---|---|---|---|---|---|
| A | | 18 | 19 | 42 | 40 | 35 | 27 | 49 | 40 | 30 | 20 | 52 | 51 | 54 | 43 | 32 | 72 | 30 | 28 | 13 | 46 | 51 | 51 | 32 | 37 | 21 | 933 | |
| G | | 22 | 26 | 31 | 33 | 38 | 46 | 24 | 33 | 43 | 53 | 21 | 22 | 19 | 30 | 41 | 1 | 43 | 45 | 31 | 27 | 22 | 22 | 41 | 36 | 52 | | 802 |
| T | 16 | 14 | 17 | | | | | | | | | | | | | | | | | 14 | | | | | | | 61 | |
| C | 57 | 19 | 11 | | | | | | | | | | | | | | | | | 15 | | | | | | | | 102 |
| **CSB424** | GGA | AGA | AGT | CTT | CTA | CTT | CTT | CTT | CTT | CTT | CTT | CTT | CTT | CTA | CTT | CTT | CTT | CTA | CTA | AGC | CTT | CTA | CTT | CTT | CTT | CTT | | |
| **CSB430** | GGT | CGT | TCT | TTA | TTA | TTA | TTA | TTA | TTA | TTA | TTA | TTA | TTA | TTA | TTA | TTA | TTA | TTA | TTA | TCA | TTA | TTA | TTA | TTA | TTA | TTA | 26 | 0 |
| **CSB426.123** | GGT | CGC | TCG | TTA | TTA | TTA | TTA | TTG | TTA | TTA | TTG | TTA | TTA | TTA | TTA | TTG | TTA | TTA | TTA | TCA | TTA | TTA | TTA | TTA | TTA | TTA | 21 | 5 |
| CSB426.111 | GGC | CGG | TCA | TTA | TTA | TTA | TTA | TTA | TTA | TTA | TTG | TTA | TTA | TTA | TTA | TTA | TTA | TTA | TTG | TCG | TTA | TTA | TTA | TTA | TTG | TTG | 19 | 7 |
| **CSB426.132** | GGT | CGT | TCG | TTA | TTA | TTG | TTA | TTA | TTA | TTA | TTA | TTA | TTA | TTA | TTG | TTG | TTA | TTG | TTA | TCT | TTA | TTG | TTA | TTA | TTA | TTG | 19 | 7 |
| **CSB426.137** | GGT | CGA | TCG | TTG | TTA | TTG | TTG | TTA | TTG | TTA | TTA | TTA | TTA | TTA | TTA | TTA | TTA | TTA | TTG | TCC | TTA | TTA | TTA | TTG | TTA | TTA | 18 | 8 |
| **CSB426.154** | GGT | CGG | TCT | TTG | TTA | TTA | TTA | TTG | TTA | TTA | TTA | TTA | TTG | TTG | TTA | TTA | TTA | TTG | TTG | TCC | TTA | TTA | TTA | TTA | TTA | TTA | 18 | 8 |
| CSB426.157 | GGC | CGT | TCA | TTA | TTA | TTA | TTA | TTA | TTG | TTG | TTG | TTA | TTA | TTA | TTG | TTG | TTA | TTG | TTA | TCT | TTA | TTA | TTA | TTG | TTA | TTG | 17 | 9 |
| CSB426.26 | GGC | CGT | TCC | TTA | TTA | TTA | TTA | TTA | TTG | TTA | TTG | TTA | TTA | TTA | TTG | TTA | TTA | TTG | TTG | TCG | TTA | TTA | TTA | TTA | TTA | TTG | 17 | 9 |
| CSB426.216 | GGC | CGG | TCT | TTA | TTA | TTA | TTA | TTG | TTG | TTG | TTA | TTA | TTG | TTA | TTA | TTA | TTA | TTA | TTA | TCA | TTA | TTA | TTA | TTG | TTG | TTG | 17 | 9 |
| CSB426.312 | GGT | CGT | TCT | TTA | TTA | TTG | TTA | TTG | TTA | TTA | TTG | TTA | TTA | TTA | TTA | TTA | TTA | TTG | TTG | TCG | TTG | TTG | TTA | TTA | TTG | TTA | 17 | 9 |
| CSB426.327 | GGC | CGA | TCA | TTA | TTA | TTA | TTA | TTA | TTG | TTA | TTG | TTA | TTA | TTA | TTA | TTA | TTA | TTG | TTG | TCC | TTG | TTA | TTA | TTG | TTG | TTA | 17 | 9 |
| CSB426.17 | GGC | CGG | TCA | TTA | TTG | TTG | TTG | TTA | TTG | TTG | TTG | TTA | TTA | TTA | TTA | TTA | TTA | TTA | TTA | TCA | TTA | TTG | TTA | TTA | TTA | TTG | 16 | 10 |
| CSB426.125 | GGC | CGA | TCC | TTA | TTA | TTA | TTG | TTA | TTA | TTA | TTG | TTA | TTA | TTA | TTG | TTG | TTA | TTG | TTA | TCG | TTA | TTA | TTA | TTA | TTG | TTG | 16 | 10 |
| CSB426.141 | GGC | CGT | TCG | TTG | TTG | TTA | TTG | TTA | TTA | TTG | TTA | TTG | TTA | TTA | TTG | TTG | TTA | TTG | TTA | TCA | TTA | TTA | TTA | TTA | TTA | TTA | 16 | 10 |
| CSB426.155 | GGT | CGT | TCC | TTA | TTA | TTA | TTG | TTA | TTG | TTA | TTG | TTA | TTA | TTA | TTG | TTG | TTA | TTA | TTG | TCT | TTG | TTA | TTA | TTA | TTG | TTG | 16 | 10 |
| CSB426.21 | GGT | CGG | TCA | TTA | TTG | TTG | TTG | TTA | TTG | TTG | TTG | TTA | TTA | TTA | TTG | TTA | TTA | TTG | TTA | TCA | TTG | TTA | TTA | TTA | TTA | TTA | 16 | 10 |
| CSB426.27 | GGC | CGC | TCG | TTG | TTA | TTA | TTA | TTA | TTA | TTG | TTG | TTA | TTG | TTA | TTA | TTA | TTA | TTA | TTA | TCA | TTA | TTG | TTG | TTA | TTA | TTG | 16 | 10 |
| CSB426.28 | GGT | CGG | TCA | TTA | TTA | TTA | TTG | TTA | TTA | TTG | TTG | TTA | TTA | TTA | TTG | TTA | TTA | TTG | TTG | TCT | TTA | TTA | TTG | TTG | TTG | TTA | 16 | 10 |
| CSB426.328 | GGC | CGA | TCT | TTA | TTA | TTG | TTG | TTA | TTA | TTG | TTG | TTA | TTA | TTA | TTA | TTG | TTA | TTG | TTG | TCG | TTA | TTA | TTA | TTA | TTA | TTG | 16 | 10 |
| CSB426.14 | GGC | CGA | TCA | TTA | TTA | TTG | TTG | TTG | TTG | TTA | TTA | TTG | TTA | TTA | TTA | TTG | TTA | TTG | TTG | TCG | TTA | TTA | TTA | TTA | TTA | TTG | 15 | 11 |
| CSB426.136 | GGC | CGG | TCA | TTA | TTA | TTG | TTG | TTA | TTA | TTG | TTG | TTA | TTA | TTA | TTA | TTG | TTA | TTA | TTA | TCC | TTG | TTA | TTA | TTG | TTA | TTG | 15 | 11 |
| CSB426.142 | GGC | CGA | TCA | TTA | TTA | TTA | TTG | TTG | TTA | TTA | TTG | TTA | TTG | TTG | TTA | TTA | TTA | TTG | TTG | TCC | TTA | TTA | TTA | TTG | TTG | TTA | 15 | 11 |
| CSB426.162 | GGT | CGC | TCT | TTG | TTG | TTA | TTA | TTA | TTG | TTA | TTG | TTA | TTA | TTG | TTA | TTG | TTA | TTG | TTG | TCT | TTA | TTA | TTA | TTG | TTA | TTG | 15 | 11 |
| CSB426.212 | GGC | CGC | TCT | TTA | TTA | TTG | TTG | TTA | TTA | TTG | TTA | TTG | TTA | TTA | TTA | TTG | TTA | TTG | TTG | TCG | TTA | TTA | TTA | TTA | TTA | TTG | 15 | 11 |
| CSB426.218 | GGC | CGC | TCC | TTA | TTG | TTA | TTA | TTA | TTG | TTA | TTA | TTA | TTA | TTA | TTA | TTG | TTA | TTG | TTG | TCA | TTA | TTA | TTA | TTG | TTG | TTG | 15 | 11 |
| CSB426.222 | GGT | CGA | TCC | TTA | TTG | TTA | TTG | TTG | TTG | TTG | TTG | TTG | TTA | TTG | TTA | TTA | TTA | TTG | TTA | TCT | TTA | TTA | TTA | TTG | TTA | TTA | 15 | 11 |
| CSB426.33 | GGC | CGT | TCT | TTA | TTG | TTA | TTA | TTA | TTA | TTG | TTG | TTG | TTA | TTA | TTG | TTG | TTA | TTA | TTG | TCA | TTA | TTG | TTA | TTA | TTG | TTG | 15 | 11 |
| CSB426.35 | GGC | CGA | TCG | TTG | TTA | TTG | TTA | TTA | TTA | TTA | TTG | TTA | TTA | TTA | TTA | TTG | TTA | TTG | TTG | TCT | TTA | TTA | TTG | TTG | TTA | TTG | 15 | 11 |
| CSB426.36 | GGC | CGC | TCC | TTA | TTA | TTA | TTG | TTA | TTG | TTG | TTG | TTA | TTA | TTA | TTA | TTA | TTA | TTA | TTA | TCG | TTA | TTG | TTA | TTA | TTG | TTG | 15 | 11 |
| CSB426.321 | GGC | CGA | TCC | TTA | TTA | TTA | TTG | TTA | TTA | TTG | TTG | TTA | TTG | TTA | TTG | TTA | TTA | TTA | TTG | TCG | TTA | TTA | TTG | TTG | TTA | TTA | 15 | 11 |
| **CSB426.324** | GGC | CGA | TCA | TTA | TTA | TTA | TTA | TTA | TTA | TTG | TTA | TTA | TTG | TTA | TTA | TTG | TTA | TTA | TTA | TCG | TTG | TTG | TTG | TTG | TTG | TTG | 15 | 11 |
| CSB426.133 | GGC | CGG | TCA | TTG | TTA | TTA | TTG | TTG | TTG | TTG | TTA | TTA | TTA | TTA | TTA | TTG | TTA | TTG | TTG | TCG | TTA | TTA | TTA | TTA | TTA | TTG | 14 | 12 |
| CSB426.214 | GGT | CGA | TCG | TTA | TTG | TTG | TTG | TTA | TTG | TTA | TTG | TTA | TTA | TTA | TTG | TTA | TTA | TTG | TTG | TCG | TTG | TTA | TTA | TTA | TTG | TTA | 14 | 12 |
| CSB426.221 | GGC | CGA | TCA | TTA | TTA | TTG | TTG | TTA | TTA | TTG | TTG | TTA | TTA | TTG | TTA | TTA | TTA | TTG | TTG | TCG | TTA | TTA | TTG | TTG | TTG | TTA | 14 | 12 |
| CSB426.31 | GGC | CGA | TCG | TTG | TTA | TTA | TTA | TTA | TTA | TTG | TTG | TTG | TTG | TTG | TTA | TTA | TTA | TTA | TTA | TCC | TTA | TTA | TTG | TTG | TTA | TTG | 14 | 12 |
| CSB426.32 | GGT | CGC | TCG | TTA | TTA | TTG | TTG | TTA | TTG | TTA | TTG | TTA | TTA | TTA | TTA | TTA | TTA | TTG | TTA | TCG | TTG | TTA | TTG | TTA | TTG | TTG | 14 | 12 |
| CSB426.37 | GGC | CGC | TCA | TTG | TTA | TTA | TTA | TTA | TTA | TTG | TTA | TTG | TTA | TTA | TTG | TTA | TTA | TTA | TTG | TCG | TTG | TTA | TTA | TTG | TTG | TTG | 14 | 12 |
| CSB426.15 | GGC | CGC | TCA | TTA | TTG | TTG | TTA | TTA | TTG | TTG | TTG | TTG | TTG | TTA | TTA | TTG | TTA | TTA | TTA | TCC | TTG | TTA | TTA | TTA | TTA | TTG | 13 | 13 |
| **CSB426.127** | GGC | CGG | TCT | TTG | TTG | TTG | TTG | TTG | TTA | TTG | TTG | TTA | TTA | TTA | TTA | TTG | TTA | TTA | TTA | TCT | TTA | TTA | TTG | TTG | TTA | TTG | 13 | 13 |
| CSB426.134 | GGC | CGC | TCA | TTG | TTG | TTG | TTA | TTA | TTG | TTA | TTG | TTA | TTA | TTA | TTG | TTA | TTA | TTG | TTG | TCT | TTA | TTG | TTA | TTG | TTA | TTG | 13 | 13 |
| CSB426.145 | GGC | CGC | TCG | TTA | TTA | TTA | TTG | TTA | TTA | TTG | TTG | TTA | TTA | TTA | TTG | TTA | TTA | TTG | TTG | TCA | TTG | TTA | TTG | TTA | TTG | TTG | 13 | 13 |
| CSB426.147 | GGC | CGT | TCA | TTA | TTA | TTA | TTG | TTA | TTA | TTG | TTG | TTG | TTG | TTA | TTA | TTG | TTA | TTA | TTG | TCG | TTG | TTG | TTA | TTA | TTG | TTG | 13 | 13 |
| CSB426.151 | GGT | CGA | TCG | TTG | TTG | TTG | TTG | TTG | TTA | TTA | TTG | TTA | TTA | TTA | TTA | TTA | TTA | TTG | TTG | TCT | TTG | TTA | TTG | TTG | TTA | TTG | 13 | 13 |
| **CSB426.156** | GGC | CGG | TCT | TTG | TTG | TTA | TTA | TTG | TTA | TTG | TTA | TTA | TTG | TTA | TTA | TTA | TTA | TTA | TTG | TCG | TTA | TTG | TTG | TTA | TTG | TTG | 13 | 13 |
| CSB426.163 | GGC | CGA | TCG | TTG | TTG | TTA | TTG | TTG | TTA | TTA | TTG | TTG | TTA | TTA | TTG | TTG | TTA | TTA | TTG | TCG | TTA | TTA | TTA | TTA | TTA | TTG | 13 | 13 |
| CSB426.215 | GGT | CGC | TCG | TTA | TTG | TTG | TTG | TTG | TTA | TTA | TTG | TTA | TTA | TTA | TTA | TTG | TTA | TTG | TTG | TCG | TTA | TTA | TTA | TTG | TTG | TTA | 13 | 13 |
| CSB426.38 | GGC | CGT | TCT | TTA | TTA | TTG | TTG | TTA | TTA | TTA | TTG | TTA | TTG | TTG | TTG | TTA | TTA | TTG | TTG | TCT | TTG | TTG | TTA | TTA | TTG | TTG | 13 | 13 |
| CSB426.313 | GGC | CGG | TCT | TTA | TTA | TTG | TTA | TTA | TTG | TTA | TTG | TTA | TTG | TTG | TTA | TTG | TTA | TTA | TTG | TCG | TTA | TTA | TTA | TTG | TTG | TTG | 13 | 13 |
| CSB426.322 | GGC | CGC | TCG | TTA | TTG | TTG | TTG | TTA | TTA | TTG | TTA | TTG | TTG | TTG | TTA | TTG | TTA | TTA | TTA | TCT | TTG | TTA | TTA | TTA | TTA | TTG | 13 | 13 |
| CSB426.334 | GGC | CGG | TCA | TTG | TTA | TTA | TTA | TTA | TTG | TTG | TTG | TTA | TTA | TTA | TTG | TTG | TTA | TTG | TTG | TCC | TTA | TTG | TTA | TTG | TTA | TTA | 13 | 13 |
| CSB426.138 | GGC | CGC | TCA | TTA | TTA | TTG | TTG | TTA | TTG | TTG | TTG | TTA | TTA | TTA | TTA | TTG | TTA | TTG | TTA | TCG | TTA | TTA | TTG | TTG | TTG | TTG | 12 | 14 |
| CSB426.144 | GGC | CGG | TCG | TTG | TTG | TTG | TTG | TTG | TTA | TTA | TTA | TTA | TTA | TTA | TTA | TTG | TTA | TTA | TTA | TCG | TTA | TTG | TTG | TTG | TTA | TTG | 12 | 14 |
| CSB426.161 | GGC | CGG | TCT | TTG | TTA | TTA | TTG | TTG | TTG | TTG | TTG | TTA | TTA | TTA | TTA | TTA | TTA | TTG | TTG | TCC | TTA | TTA | TTG | TTG | TTA | TTG | 12 | 14 |
| CSB426.23 | GGC | CGT | TCG | TTG | TTG | TTG | TTG | TTA | TTA | TTG | TTG | TTA | TTA | TTA | TTG | TTG | TTA | TTG | TTA | TCC | TTG | TTA | TTA | TTA | TTA | TTG | 12 | 14 |
| **CSB426.323** | GGC | CGG | TCT | TTG | TTA | TTG | TTG | TTA | TTG | TTG | TTG | TTG | TTA | TTA | TTA | TTA | TTA | TTG | TTG | TCA | TTG | TTA | TTA | TTG | TTA | TTG | 12 | 14 |
| CSB426.124 | GGC | CGC | TCT | TTG | TTG | TTG | TTG | TTA | TTG | TTA | TTG | TTG | TTA | TTA | TTA | TTG | TTA | TTA | TTA | TCC | TTA | TTG | TTA | TTG | TTG | TTG | 11 | 15 |
| CSB426.126 | GGC | CGG | TCG | TTG | TTA | TTG | TTG | TTG | TTG | TTG | TTA | TTA | TTA | TTA | TTG | TTG | TTA | TTA | TTA | TCC | TTA | TTG | TTA | TTA | TTG | TTG | 11 | 15 |
| CSB426.128 | GGC | CGG | TCG | TTG | TTG | TTG | TTA | TTA | TTG | TTG | TTG | TTA | TTA | TTA | TTG | TTA | TTA | TTA | TTG | TCC | TTA | TTA | TTG | TTG | TTG | TTA | 11 | 15 |
| CSB426.135 | GGT | CGA | TCG | TTG | TTG | TTG | TTG | TTG | TTG | TTA | TTA | TTA | TTA | TTG | TTA | TTG | TTA | TTA | TTG | TCG | TTG | TTA | TTG | TTG | TTA | TTG | 11 | 15 |
| CSB426.211 | GGC | CGT | TCG | TTG | TTG | TTG | TTA | TTA | TTA | TTG | TTG | TTA | TTA | TTG | TTG | TTG | TTA | TTA | TTG | TCA | TTG | TTA | TTA | TTG | TTG | TTG | 11 | 15 |
| CSB426.333 | GGC | CGG | TCG | TTG | TTA | TTG | TTA | TTA | TTA | TTG | TTG | TTA | TTG | TTG | TTG | TTA | TTA | TTG | TTG | TCG | TTA | TTA | TTG | TTG | TTG | TTA | 11 | 15 |
| CSB426.335 | GGT | CGG | TCG | TTG | TTG | TTA | TTG | TTA | TTG | TTG | TTA | TTG | TTG | TTG | TTA | TTG | TTA | TTG | TTA | TCT | TTA | TTA | TTA | TTG | TTG | TTG | 11 | 15 |
| CSB426.12 | GGC | CGG | TCG | TTG | TTG | TTG | TTG | TTA | TTA | TTG | TTG | TTG | TTA | TTG | TTG | TTG | TTA | TTA | TTA | TCA | TTG | TTG | TTA | TTA | TTG | TTA | 10 | 16 |
| CSB426.131 | GGC | CGG | TCG | TTG | TTG | TTG | TTA | TTA | TTG | TTG | TTA | TTG | TTA | TTA | TTG | TTA | TTA | TTG | TTG | TCG | TTG | TTA | TTA | TTG | TTA | TTG | 10 | 16 |
| CSB426.143 | GGC | CGA | TCT | TTA | TTA | TTG | TTG | TTA | TTA | TTG | TTA | TTG | TTG | TTG | TTG | TTG | TTA | TTG | TTG | TCG | TTG | TTA | TTG | TTG | TTA | TTG | 10 | 16 |
| **CSB426.148** | GGC | CGT | TCC | TTA | TTG | TTA | TTA | TTG | TTG | TTG | TTA | TTG | TTG | TTG | TTA | TTG | TTG | TTG | TTG | TCT | TTG | TTG | TTA | TTG | TTA | TTA | 10 | 16 |
| CSB426.164 | GGC | CGC | TCC | TTA | TTG | TTA | TTG | TTG | TTA | TTG | TTG | TTA | TTG | TTA | TTG | TTG | TTA | TTG | TTG | TCA | TTA | TTA | TTG | TTA | TTG | TTG | 10 | 16 |
| CSB426.24 | GGC | CGT | TCC | TTA | TTA | TTG | TTG | TTG | TTA | TTG | TTG | TTA | TTG | TTG | TTA | TTG | TTA | TTA | TTG | TCG | TTA | TTG | TTA | TTG | TTG | TTG | 10 | 16 |
| CSB426.217 | GGC | CGT | TCT | TTA | TTG | TTG | TTG | TTG | TTA | TTA | TTG | TTG | TTA | TTA | TTA | TTG | TTA | TTG | TTG | TCC | TTG | TTG | TTG | TTA | TTG | TTG | 10 | 16 |
| CSB426.34 | GGC | CGA | TCG | TTG | TTG | TTG | TTG | TTG | TTA | TTG | TTG | TTA | TTG | TTG | TTG | TTA | TTA | TTA | TTA | TCG | TTA | TTG | TTG | TTG | TTA | TTA | 10 | 16 |
| **CSB426.331** | GGC | CGG | TCG | TTG | TTG | TTA | TTG | TTA | TTA | TTA | TTG | TTA | TTA | TTA | TTG | TTG | TTA | TTG | TTG | TCG | TTA | TTA | TTG | TTG | TTG | TTG | 10 | 16 |
| **CSB426.326** | GGC | CGC | TCA | TTA | TTG | TTA | TTG | TTG | TTG | TTG | TTG | TTG | TTA | TTA | TTA | TTG | TTA | TTG | TTA | TCG | TTG | TTG | TTA | TTG | TTG | TTG | 9 | 17 |
| **CSB426.332** | GGC | CGC | TCT | TTG | TTG | TTG | TTG | TTG | TTG | TTA | TTG | TTA | TTG | TTA | TTG | TTG | TTA | TTG | TTG | TCC | TTG | TTA | TTA | TTG | TTA | TTA | 9 | 17 |
| **CSB426.158** | GGC | CGC | TCC | TTA | TTG | TTA | TTG | TTA | TTG | TTA | TTG | TTG | TTG | TTG | TTG | TTA | TTA | TTG | TTA | TCG | TTG | TTG | TTA | TTG | TTG | TTG | 8 | 18 |

**Fig. S13. Summary of the codon compressed *rbcL* coding sequences.** Composition of the 26 target codons in 73 transformants selected from the CSB426 library are compared to those in the WT *rbcL* (CSB424) and the *rbcL* compressed using the defined compression scheme (CSB430). Sequences are sorted based on the decreasing number of codons ending with A or T. Randomised codons are coloured by the third base of the codon: T - red, C - blue, A - green, G - black. Counts of codons are presented in at the top of the figure. Sequences highlighted in yellow were selected for subsequent analysis.

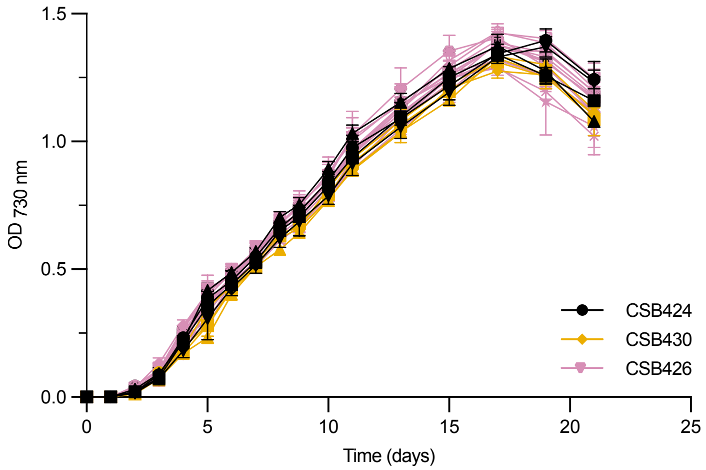


**Fig. S13. Phototrophic growth of the *rbcL* complemented lines.** Selected transformants of pCSB424 (black; n=4), pCSB430 (orange; n=4) and pCSB426 (pink; n=13) were grown in HSM at light intensity of 80 µmol·m$^{-2}$·s$^{-1}$. $OD_{730nm}$ of cultures was measured every 24-48 h. Error bars represent standard deviations for three independent cultures.

**Table S1.** List of plasmids generated and used in this study. For level 2 plasmids assembled by Golden Gate cloning, individual level 1 plasmids are highlighted with alternating highlighting. Sequences are available in the GenBank database (www.ncbi.nlm.nih.gov/genbank/) with the appropriate accession numbers.

| Plasmid | Backbone | Level | Description | Accession number |
|---|---|---|---|---|
| pPM900 | - | - | Level 2 recipient plasmid (XY) | MT361981 |
| pCSB19 | - | - | Level 2 recipient plasmid (AZ) | PP746763 |
| pCSB245 | pPM900 | 2 | LHA(rpoA)-PatpA-5'atpA-codA.aadA-TrbcL-PpsbD-5'psbD-rpoA_FLAG(WT)-RHA(rpoA) | PV101188 |
| pCSB246 | pPM900 | 2 | LHA(rpoA)-PatpA-5'atpA-codA.aadA-TrbcL-PpsbD-5'psbD-rpoA_FLAG(Def.comp.)-RHA(rpoA) | PV101189 |
| pCSB247 | pPM900 | 2 | LHA(rpoA)-PatpA-5'atpA-codA.aadA-TrbcL-PpsbD-5'psbD-rpoA_FLAG(ChimeraMAP)-RHA(rpoA) | PV101190 |
| pCSB248 | pPM900 | 2 | LHA(rpoA)-PatpA-5'atpA-codA.aadA-TrbcL-PpsbD-5'psbD-rpoA_FLAG(CUO)-RHA(rpoA) | PV101191 |
| pCSB317 | pPM900 | 2 | LHA(ycf1)-PatpA-5'atpA-codA.aadA-TrbcL-PpsbD-5'psbD-ycf1(WT)-RHA(ycf1) | PV101193 |
| pCSB322 | pPM900 | 2 | LHA(rpoA)-PpsaA-5'psaA-aphA6-TatpB-PrpoA-5'rpoA-rpoA_FLAG(Def.comp.) | PV101192 |
| pCSB323 | pPM900 | 2 | LHA(psbA)-PatpA-5'atpA-codA.aadA-TrbcL-RHA(psbA) | PV101195 |
| pCSB347 | pPM900 | 2 | LHA(rpoB2)-PpsaA-5'psaA-aphA6-TatpB-psbE-rps2 operon(Def.comp)-PrpoA-5'rpoA-rpoA(Def.comp) | PV101200 |
| pCSB416 | pPM900 | 2 | LHA(ycf1)-PatpA-5'atpA-codA.aadA-TrbcL-PpsbD-5'psbD-ycf1(Def.comp.)-RHA(ycf1) | PV101194 |
| pCSB419 | pCSB19 | 2 | LHA(rbcL)-PrbcL-5'rbcL-codA.aadA-TrbcL-RHA(rbcL) | PV101201 |
| pCSB424 | pCSB19 | 2 | LHA(rbcL)-PrbcL-5'rbcL-rbcL(WT)-TrbcL-RHA(rbcL) | PV101202 |
| pCSB430 | pCSB19 | 2 | LHA(rbcL)-PrbcL-5'rbcL-rbcL(Def.comp.)-TrbcL-RHA(rbcL) | PV101203 |
| pCSB426 library | pCSB19 | 2 | LHA(rbcL)-PrbcL-5'rbcL-rbcL(Combinatorial library)-TrbcL-RHA(rbcL) | PV101204 |
| pCSB487 | pCSB19 | 2 | LHA(psbA)-PpsbA-psbA-TpsbA-RHA(psbA) | PV101196 |
| pCSB488 | pCSB19 | 2 | LHA(psbA)-PpsbA-psbA(no-introns)-TpsbA-RHA(psbA) | PV101197 |
| pCSB489 | pCSB19 | 2 | LHA(psbA)-PpsbA-psbA(no-introns,Def.comp.)-TpsbA-RHA(psbA) | PV101198 |
| pCSB491 | pPM900 | 2 | LHA(psaA.ex3)-PatpA-5'atpA-codA.aadA-TrbcL-PpsaA.ex1-5'psaA.ex1-psaA.ex1+2+3(Def.comp.)-RHA(TpsaA.ex3) | PV101199 |

**Table S2.** List of *Chlamydomonas reinhardtii* strains used or generated in this study. Transgenic strains from this study were generated by the introduction of the relevant plasmids detailed in Table S1.

| Strain | Genetic background | Description | Reference |
|---|---|---|---|
| CC-1690 | - | WT strain (21 gr) | (7) |
| CSB128 (Luc1.R2) | CC-1690 | WT CC-1690 containing *lucCP* firefly luciferase cassette in the *rrn5-psbA* insertion site | (8) |
| CSB128-358 | CSB128 | Transformed with plasmid pCSB358. Inverted repeat replaced with *aphA6* cassette. | This work |
| CSB245<br>CSB246<br>CSB247<br>CSB248 | CSB128 | Transformed with plasmids pCSB245 (WT *rpoA* CDS), pCSB246 (Def. comp.), pCSB247 (ChimeraMAP) and pCSB248 (CUO), resulting in recoded rpoA CDS, *rpoA* promoter and 5'UTR swapped for those of *psbD,* deletion of SDRs, and integration of the FLAG-tag | This work |
| CSB322 | CSB246 | CSB246-1 transformed with pCSB322. *rpoA* promoter and 5'UTR restored upstream of *rpoA* CDS | This work |
| CSB317<br>CSB416 | CS128 | Transformed with plasmids pCSB317 (WT *ycf1* CDS) and pCSB416 (*ycf1* Def. comp.), resulting in recoded *ycf1* CDS, *ycf1* promoter and 5'UTR swapped for those of *psbD* | This work |
| CSB323 | CSB128-358 | Transformed with plasmid pCSB323, *psbA* gene replaced with the *codA.aadA* cassette. Non-photosynthetic phenotype. | This work |
| CSB487<br>CSB488<br>CSB489 | CSB323 | *psbA* knockout complemented with pCSB487 (*psbA* WT), pCSB488 (*psbA* WT, no introns) or pCSB489 (*psbA* Def. comp., no introns) | This work |
| CSB491 | CSB128 | Transformed with pCSB491, *psaA* made intron-less and recoded using Def. comp. scheme | This work |
| CSB347 | CSB246 | CSB246-1 transformed with pCSB347. *trnM$^{e}$-rps2* operon recoded according to the Def. comp. scheme. | This work |
| CSB419 | CC-1690 | Transformed with plasmid pCSB419, *rbcL* CDS replaced with *codA.aadA*. Non-photosynthetic and light sensitive phenotype. | This work |
| CSB424<br>CSB426<br>CSB430 | CSB419 | *rbcL* knockout complemented with pCSB424 (*rbcL* WT), pCSB430 (*rbcL* Def. comp.) or pCSB426 library (combinatorial assembly of codons for target positions) | This work |

**Table S3.** List of synthetic oligonucleotides used in this study.

| Name | Sequence | Description |
|---|---|---|
| **Genotyping of CSB245, CSB246, CSB247 and CSB248 transformants** | | |
| oligoCSB534 | TGGTTTCAATTATTAAAGGTCAAGG | *rpoA* locus F |
| oligoCSB535 | CTCCCTCGTACAAAGCATAC | *rpoA* locus R |
| oligoCSB536 | CGTGTTGAATACCGTTAGCA | *codA.aadA* cassette R |
| **Amplification of *rpoA* CDS in CSB246 and CSB248, and Sanger sequencing** | | |
| oligoCSB145 | GCGCAGATCAGTTGGAAGA | *rpoA* amplification F |
| oligoCSB459 | TGAAGGTTTTTGGTGGAATTG | *rpoA* sequencing |
| oligoCSB471 | TCAACAAATATGAATTATCGAGCAA | *rpoA* sequencing |
| oligoCSB501 | TTTTAGTTGCTCAAAGGGGTTT | *rpoA* sequencing |
| oligoCSB537 | TGCCAGTTTCCTCCTTTCTC | *rpoA* amplification R |
| **Genotyping of CSB322 transformants** | | |
| oligoCSB95 | GTTCTTAGAGCTAAAAGAGAAGAACAA | *codA.aadA* cassette R |
| oligoCSB534 | TGGTTTCAATTATTAAAGGTCAAGG | *rpoA* locus F |
| oligoCSB535 | CTCCCTCGTACAAAGCATAC | *rpoA* locus R |
| oligoCSB611 | CTTCACGAGAAACTGAGTATG | *aphA6* cassette R |
| **Amplification of *rpoA* in CSB322 for Amplicon-NGS** | | |
| oligoCSB713 | ACACTCTTTCCCTACACGACGCTCTTCCGATCTATGAC AATTTATCCAAATTTAAAAAAAATC | *rpoA* product 1 F |
| oligoCSB714 | GACTGGAGTTCAGACGTGTGCTCTTCCGATCTATTTCA TATTTAAAAAACCATCCTC | *rpoA* product 1 R |
| oligoCSB715 | ACACTCTTTCCCTACACGACGCTCTTCCGATCTCAACA TTAGCTGAGGATGG | *rpoA* product 2 F |
| oligoCSB716 | GACTGGAGTTCAGACGTGTGCTCTTCCGATCTGCTTTG TTAACGAATTTATGC | *rpoA* product 2 R |
| oligoCSB717 | ACACTCTTTCCCTACACGACGCTCTTCCGATCTCTGAC CCATCTTTAGAGTTTTC | *rpoA* product 3 F |
| oligoCSB718 | GACTGGAGTTCAGACGTGTGCTCTTCCGATCTCCTAAA AAAGTTTTAATTTTTGTTAATG | *rpoA* product 3 R |
| oligoCSB719 | ACACTCTTTCCCTACACGACGCTCTTCCGATCTTCAAC AGGCATTTCAAATACTG | *rpoA* product 4 F |
| oligoCSB720 | GACTGGAGTTCAGACGTGTGCTCTTCCGATCTAGATTA ATGCTTGATATAAAGCTTTG | *rpoA* product 4 R |
| oligoCSB721 | ACACTCTTTCCCTACACGACGCTCTTCCGATCTTCCAC GCAAAGCTTTATATC | *rpoA* product 5 F |
| oligoCSB722 | GACTGGAGTTCAGACGTGTGCTCTTCCGATCTATCAGT AACTAAAGTTAAGCCTAAC | *rpoA* product 5 R |
| **Genotyping of CSB317 and CSB416 transformants** | | |
| oligoCSB95 | GTTCTTAGAGCTAAAAGAGAAGAACAA | *codA.aadA* cassette R |
| oligoCSB219 | TCAACAGCACCTGTAATTGCTA | *ycf1* locus F |
| oligoCSB711 | TAGCTTGTCACTGCCATTCC | *ycf1* promoter F |
| oligoCSB712 | ACTGCAAACGCAAAACATTG | *ycf1* promoter R |
| **Amplification of ycf1 in CSB416 and Amplicon-NGS** | | |
| oligoCSB921 | ATTCAATTGAAGAAGATGATTTAC | *ycf1* product d F |
| oligoCSB922 | CTTGTACCACTTTAGTTAATTG | *ycf1* product d R |
| oligoCSB923 | AGCTACACGTAAACCACG | *ycf1* product c F |
| oligoCSB924 | GTATTTGATAATGGTTTAAATGAAAC | *ycf1* product c R |
| oligoCSB929 | GAATTTGGTGCGGTTTTTAC | *ycf1* product a F |
| oligoCSB930 | CCATGGGATTACAAAAAAACG | *ycf1* product a R |
| oligoCSB931 | GTAGCTTCATTACGTCGTG | *ycf1* product b F |
| oligoCSB932 | TGTATCATAATGTTCCGATAATAAG | *ycf1* product b R |
| **Genotyping of CSB323 transformants** | | |

| | | |
|---|---|---|
| oligoCSB37 | CAATGACAGAAAAAGAAATTGTTGA | *lucCP* CDS R |
| oligoCSB95 | GTTCTTAGAGCTAAAAGAGAAGAACAA | *codA.aadA* CDS R |
| oligoCSB591 | GGTAGGTTCTGTCACTGAC | *psaA* 3'UTR F |
| oligoCSB728 | GGGAAGAGAATGGGTTCGAT | *psbA* left flank F |
| oligoCSB731 | GGCTAGAATTTTCACGACGTTC | *psbA ex 1* CDS R |
| **Genotyping of CSB487, CSB488 and CSB489 transformants** | | |
| oligoCSB95 | GTTCTTAGAGCTAAAAGAGAAGAACAA | *codA.aadA* CDS R |
| oligoCSB728 | GGGAAGAGAATGGGTTCGAT | *psbA* left flank F |
| oligoCSB1020 | TGGTAAGGACCACCGTTGTA | *psbA ex 2* CDS R |
| **Genotyping of CSB491 transformants** | | |
| oligoCSB95 | GTTCTTAGAGCTAAAAGAGAAGAACAA | *codA.aadA* CDS R |
| oligoCSB1104 | TAACCTGTCGCAGGCCATTA | *wendy2* F |
| oligoCSB1105 | CCTTGGAAACCACCACCTAC | *psaA ex 3* CDS R |
| oligoCSB1106 | CGGCACGTAGTTGGAAAGTA | *psaA ex 3* g. II intron F |
| **Genotyping of CSB347 transformants** | | |
| oligoCSB95 | GTTCTTAGAGCTAAAAGAGAAGAACAA | *codA.aadA* CDS R |
| oligoCSB534 | TGGTTTCAATTATTAAAGGTCAAGG | *rpoA* locus F |
| oligoCSB535 | CTCCCTCGTACAAAGCATAC | *rpoA* locus R |
| oligoCSB611 | CTTCACGAGAAACTGAGTATG | *aphA6* cassette R |
| oligoCSB629 | CGGACGACATGGGAATAAAG | *rpoB2* CDS F |
| oligoCSB738 | GCCTCTTACTCCCAGATATCCA | P*trnMe* R |
| **Amplification of *trnM*$^{e}$*-rps2* operon in CSB416 and Amplicon-NGS** | | |
| oligoCSB815 | ACACTCTTTCCCTACACGACGCTCTTCCGATCTCTATG TTAAACAAAAAGCCAC | product 1 F |
| oligoCSB816 | GACTGGAGTTCAGACGTGTGCTCTTCCGATCTTTTCAC CAATTGTAATAGCAG | product 1 R |
| oligoCSB817 | ACACTCTTTCCCTACACGACGCTCTTCCGATCTTAATT TAGCAGCATTAACACCAG | product 2 F |
| oligoCSB818 | GACTGGAGTTCAGACGTGTGCTCTTCCGATCTGTTAAA TTGCCTGCTGAAC | product 2 R |
| oligoCSB821 | ACACTCTTTCCCTACACGACGCTCTTCCGATCTCAAAA GTTGAAGCACGTC | product 4 F |
| oligoCSB822 | GACTGGAGTTCAGACGTGTGCTCTTCCGATCTTACCAG ACGTTGAGTTTG | product 4 R |
| oligoCSB823 | ACACTCTTTCCCTACACGACGCTCTTCCGATCTGAAAA ACGTCAAACAATTAAAG | product 5 F |
| oligoCSB824 | GACTGGAGTTCAGACGTGTGCTCTTCCGATCTAACCAT ATATAAATTTTGATTGTCTTG | product 5 R |
| oligoCSB825 | ACACTCTTTCCCTACACGACGCTCTTCCGATCTAAATT ACGTGAGTTAAAAGC | product 6 F |
| oligoCSB826 | GACTGGAGTTCAGACGTGTGCTCTTCCGATCTTTTAAT TTGACGTAATTCAGCAAC | product 6 R |
| oligoCSB827 | ACACTCTTTCCCTACACGACGCTCTTCCGATCTAGTTA AAACTAAAATGAACGTTTATG | product 7 F |
| oligoCSB828 | GACTGGAGTTCAGACGTGTGCTCTTCCGATCTCATTCA CGTACAGCATTC | product 7 R |
| oligoCSB835 | ACACTCTTTCCCTACACGACGCTCTTCCGATCTCGGTA GGTTTAGATACACG | product 3 F |
| oligoCSB836 | GACTGGAGTTCAGACGTGTGCTCTTCCGATCTTAGTTA ATTGTGCTAAAGCTTTAG | product 3 R |
| oligoCSB837 | ACACTCTTTCCCTACACGACGCTCTTCCGATCTCTAAA ATGAAATATCCTATGGATTC | product 8 F |

| | | |
|---|---|---|
| oligoCSB838 | GACTGGAGTTCAGACGTGTGCTCTTCCGATCTGACGTC CTAGAATGACAAC | product 8 R |
| oligoCSB875 | AAAAGCTAAGTAAGCACAC | product 9 F |
| oligoCSB876 | GCTAACTGACATCAATGGAC | product 9 R |
| oligoCSB877 | TAGCAAAAGATGAAAAAATCTCTCA | product 10 F |
| oligoCSB878 | CAGTTTGGCCCATTAAAC | product 10 R |
| oligoCSB879 | CACAAATTTAATGCAATTTTCAG | product 11 F |
| oligoCSB880 | GCTAACTATTACACTTCTTTGG | product 11 R |
| oligoCSB881 | AACTCAATCAATTCTATATGACAC | product 12 F |
| oligoCSB882 | CAACACGAATAGCTTCAATATC | product 12 R |
| oligoCSB883 | TTGTTCCGTTTTTCCCAC | product 13 F |
| oligoCSB884 | TAATAACATCTGTTTAGACGTTTTG | product 13 R |
| oligoCSB887 | TGTCAGCACAAGCTGAAG | product 15 F |
| oligoCSB888 | CTTAGAGAAGAGTAAAAATCAATCTG | product 15 R |
| oligoCSB895 | TTTATTTATTTATGCGCGGTTTAAAATC | product 16 F |
| oligoCSB896 | CAAGTTTTTTATAACACTGGATTTG | product 16 R |
| oligoCSB897 | TCGCTGATTTTATTTTTAATGC | product 14 F |
| oligoCSB899 | GCTTCTTTCCAGTAATCTGC | product 14 R |
| **Genotyping of CSB419** | | |
| oligoCSB376 | TGGAGTCATACGGAATGCAG | *rbcL* CSD R |
| oligoCSB536 | CGTGTTGAATACCGTTAGCA | *codA.aadA* CDS R |
| oligoCSB873 | GCGATTGCTGTTTTACCTGTT | *atpA* CDS F |
| **Generation of pCSB426 library** | | |
| oligoCSB849 | GTGTTGCGTCTCATGCGTGAGACCCACGAAGTGATCCG TTTAAAC | fragment A5 F |
| oligoCSB850 | CAGTGTCGTCTCAATGTTGAGACCCAGAGTGTTCAACC CCAG | fragment A5 R |
| oligoCSB851 | GTGTTGCGTCTCAACATACTACACACCTGATTACGTAG TACGNGATACTGATATTTTAGCTGCATTC | fragment A1 F |
| oligoCSB852 | CAGTGTCGTCTCAAACGGTCYAANGATGTTAAACCGTC AGTCCATAC | fragment A1 R |
| oligoCSB853 | GTGTTGCGTCTCACGTTACAAAGGTCGTTGTTAC | fragment A2 F |
| oligoCSB854 | CAGTGTCGTCTCACAGGTGGAATACGYAAGTCTTCYAA ACGYAAAGCACGTAAAGCTTTGAAAC | fragment A2 R |
| oligoCSB855 | GTGTTGCGTCTCACCTGCTTACGTTAAAACATTC | fragment A3 F |
| oligoCSB856 | CAGTGTCGTCTCATTGATTGTACAACCTAAYAAACCAC GACCATATTTGTTTAATTTG | fragment A3 R |
| oligoCSB857 | TCAAACCTAAATTAGGTTTRTCAGCTAAAAACTACGGT CGTGCAGTTTATGAATGTTTACGTGGTGGTTTRGACTT TACTAAAGACGACGAAAACGTAAACTCACAACCATTCA | fragment A4 sense |
| oligoCSB858 | CGCATGAATGGTTGTGAGTTTACGTTTTCGTCGTCTTT AGTAAAGTCYAAACCACCACGTAAACATTCATAAACTG CACGACCGTAGTTTTTAGCTGAYAAACCTAATTTAGGT | fragment A4 antisense |
| oligoCSB859 | AAGGGGTTGAAGACAATGCCGGTCTCATGCGTTGGCGT GACCGTTTCTTRTTCGTTGCTGAAGCTATTTAC | fragment D1 F |
| oligoCSB860 | CAGTGTTGGAAGACAAGGTCAATAACCGCGTGCATAGC ACGGTGGATGTGYAAYAAYAAACCGTTGTCACGACAG | fragment D1 R |
| oligoCSB861 | GACCGTCAACGTAACCACGGTATTCACTTCCGTGTTTT RGCTAAAGCTTTRCGTATGTCTGGTGGTGACCACTTRC ACTCTGGTACTGTTGTAGGTAAATTRGAAGGTGAACGT GAAGTTACTTTRGGTTTCGTAGACTTAATGCGTGATGA | fragment D2 sense |
| oligoCSB862 | GTAGTCATCACGCATTAAGTCTACGAAACCYAAAGTAA CTTCACGTTCACCTTCYAATTTACCTACAACAGTACCA | fragment D2 antisense |

| | | |
|---|---|---|
| | GAGTGYAAGTGGTCACCACCAGACATACGYAAAGCTTT AGCYAAAACACGGAAGTGAATACCGTGGTTACGTTGAC | |
| oligoCSB863 | AAGGGGTTGAAGACAACTACGTTGAAAAAGACCGTTCN CGTGGTATTTACTTCACTCAAGAC | fragment D3 F |
| oligoCSB914 | GTTGGAAGACAAtcccGGTCTCCGGAGCGTTACCCCAA GGGTGACCYAAAGTACCACCACCGAACTGYAAACATGC GTCATCACCGAAG | fragment D3 R |
| oligoCSB870 | CAGTGTTGGAAGACAATTCGAATTTGATACTATTGACA AATTRTAAtttttatttttcatgatgtttatgTG | fragment E1 F |
| oligoCSB915 | AAGGGGTTGAAGACAATAGTCTTTTACACCGGCTTTGA ARCCAGCACCTGCTTTAGTTTC | fragment E1 R |
| oligoCSB916 | AAGGGGTTGAAGACAAACTACCGTTTAACATTGAGACC TAGGGCAGTGAGCGCAAC | fragment E2 F |
| oligoCSB917 | AGTGTTGGAAGACAAGAGCTTGAGTACAAGCTTCYAAA GCTACACGGTTAGCTGCAGCACCTGGAGGGAGACCCTA TGCGGCATCAGAG | fragment E2 R |
| oligoCSB918 | GCTCGTAACGAAGGTCGTGACTTRGCTCGTGAAGGTGG CGACGTAATTCGTTCAGCTTGTAAATGGTCTCCAGAAT TRGCTGCTGCATGTGAAGTTTGGAAAGAAATTAAA | fragment E3 sense |
| oligoCSB919 | CGAATTTAATTTCTTTCCAAACTTCACATGCAGCAGCY AATTCTGGAGACCATTTACAAGCTGAACGAATTACGTC GCCACCTTCACGAGCYAAGTCACGACCTTCGTTAC | fragment E3 antisense |
| **Amplification of *rbcL in* CSB426 transformants and Sanger sequencing** | | |
| oligoCSB433 | CCCTGACAGGAATATACATGGTT | *rbcL* locus F |
| oligoCSB933 | GACAGGCAAATTTAAACAAAAGA | *rbcL* locus R |

**Table S4.** Efficiency of chloroplast transformations

| Construct | Unique codons | Modified codons | Modified bases | Transformants tested | Correct integration | Homoplasmy |
|---|---|---|---|---|---|---|
| **Recoding of *rpoA*** | | | | | | |
| pCSB245 (WT-FLAG) | 57 | - | - | 22 | 19 (86%) | 17 (89%) |
| pCSB246 (Def. comp.) | 47 | 56 (7.5%) | 99 (4.4%) | 16 | 14 (87%) | 14 (100%) |
| pCSB247 (ChimeraMAP) | 48 | 176 (23.5%) | 346 (15.4%) | 16 | 10 (62%) | 0 (0%) |
| pCSB248 (CUO) | 44 | 176 (23.5%) | 339 (15.1%) | 16 | 12 (75%) | 7 (58%) |
| pCSB322 (Def. comp.) | - | - | - | 16 | 15 (95%) | 15 (100%) |
| **Recoding of *ycf1*** | | | | | | |
| pCSB317 (WT) | 59 | - | - | 6 | 6 (100%) | 6 (100%) |
| pCSB416 (Def. comp.) | 50 | 147 (7.1%) | 245 (4.1%) | 18 | 18 (100%) | 14 (78%) |
| **Deletion of *psbA*** | | | | | | |
| pCSB323 | - | - | - | 22 | 19 (86%) | 4 (21%) |
| **Restoration of *psbA*** | | | | | | |
| pCSB487 (WT) | 37 | - | - | 4 | 4 (100%) | 4 (100%) |
| pCSB488 (No introns) | 37 | - | - | 6 | 6 (100%) | 5 (80%) |
| pCSB489 (Def. comp., no introns) | 34 | 16 (4.5%) | 28 (2.6%) | 3 | 3 (100%) | 3 (100%) |
| **Recoding of *psaA*** | | | | | | |
| pCSB491 | 41 | 46 (6.1%) | 91 (4%) | 24 | 20 (83%) | 20 (100%) |
| **Recoding of *trnM*$^{e}$*-psbE-rps9-ycf4-ycf3-rps18-rps2* operon** | | | | | | |
| pCSB347 | 49 | 104 (6.1%) | 174 (3.4%) | 40 | 32 (80%) | 32 (100%) |
| **Deletion of *rbcL*** | | | | | | |
| pCSB419 | - | - | - | 16 | 16 (100%) | 16 (100%) |
| **Restoration of *rbcL*** | | | | | | |
| pCSB424 | 41 | - | - | 8 | 8 (100%) | 8 (100%) |
| pCSB426 | Combinatorial library | | | 83 | 73 (88%) | 73 (100%) |
| pCSB430 | 35 | 27 (5.7%) | 48 (3.4%) | 8 | 7 (87%) | 7 (100%) |

### SI References


1. S. D. Gallaher, et al., High-throughput sequencing of the chloroplast and mitochondrion of *Chlamydomonas reinhardtii* to generate improved de novo assemblies, analyze expression patterns and transcript speciation, and evaluate diversity among laboratory strains and wild isolates. *Plant J.* 93, 545–565 (2018).
2. P. P. Chan, T. M. Lowe, tRNAscan-SE: Searching for tRNA genes in genomic Sequences. *Methods Mol. Biol.* 1962, 1–14 (2019).
3. V. Cognat, et al., PlantRNA, a database for tRNAs of photosynthetic eukaryotes. *Nucleic Acids Res.* 41, D273–9 (2013).
4. S. Alkatib, et al., The contributions of wobbling and superwobbling to the reading of the genetic code. *PLoS Genet*. 8, e1003076 (2012).
5. M. Fages-Lartaud, K. Hundvin, M. F. Hohmann-Marriott, Mechanisms governing codon usage bias and the implications for protein expression in the chloroplast of *Chlamydomonas reinhardtii*. *Plant J.* (2022). https://doi.org/10.1111/tpj.15970.
6. M. G. Kluesner, et al., EditR: A method to quantify base editing from Sanger sequencing. *CRISPR J.* 1, 239–250 (2018).
7. R. Sager, Inheritance in the green alga *Chlamydomonas reinhardi*. *Genetics* 40, 476–489 (1955).
8. H. O. Jackson, et al., CpPosNeg: A positive-negative selection strategy allowing multiple cycles of marker-free engineering of the *Chlamydomonas* plastome. *Biotechnol. J.* e2200088 (2022).
